# BNP-Track: A framework for multi-particle superresolved tracking

**DOI:** 10.1101/2023.04.03.535440

**Authors:** Lance W.Q. Xu, Ioannis Sgouralis, Zeliha Kilic, Steve Pressé

**Author notes:** Corresponding author. Email address Website: https://cbp.asu.edu/content/steve-presse-lab. Contributed equally (Lance W.Q. Xu, Ioannis Sgouralis).

## Abstract

When tracking fluorescently labeled molecules (termed “emitters”) under widefield microscopes, point spread function overlap of neighboring molecules is inevitable in both dilute and especially crowded environments. In such cases, superresolution methods leveraging rare photophysical events to distinguish static targets nearby in space introduce temporal delays that compromise tracking. As we have shown in a companion manuscript, for dynamic targets, information on neighboring fluorescent molecules is encoded as spatial intensity correlations across pixels and temporal correlations in intensity patterns across time frames. We then demonstrated how we used all spatiotemporal correlations encoded in the data to achieve superresolved tracking. That is, we showed the results of full posterior inference over both the number of emitters and their associated tracks simultaneously and self-consistently through Bayesian nonparametrics. In this companion manuscript we focus on testing the robustness of our tracking tool, BNP-Track, across sets of parameter regimes and compare BNP-Track to competing tracking methods in the spirit of a prior Nature Methods tracking competition. We explore additional features of BNP-Track including how a stochastic treatment of background yields greater accuracy in emitter number determination and how BNP-Track corrects for point spread function blur (or “aliasing”) introduced by intraframe motion in addition to propagating error originating from myriad sources (such as criss-crossing tracks, out-of-focus particles, pixelation, shot and camera artefact, stochastic background) in posterior inference over emitter numbers and their associated tracks. While head-to-head comparison with other tracking methods is not possible (as competitors cannot simultaneously learn molecule numbers and associated tracks), we can give competing methods some advantages in order to perform approximate head-to-head comparison. We show that even under such optimistic scenarios, BNP-Track is capable of tracking multiple diffraction-limited point emitters conventional tracking methods cannot resolve thereby extending the superresolution paradigm to dynamical targets.

## 1 Introduction

Engineered fluorescent tags and their photophysical response to excitation light, have been critical in localizing fluorescently labeled molecules (also referred to as “emitters”) below light’s diffraction limit [1–3] often down to a resolution of 20 nm to 30 nm within individual image frames [4–10] or better with structured excitation patterns [11]. The ability to accurately localize emitters at this scale has contributed to the visualization of T-cell antigen recognition [12], understanding the role of the plasma membrane structure in cell communication [13], and is the basis for spatial transcriptomics [14, 15] and ensuing analysis unraveling gene transcriptional models from RNA counts [16].

Despite progress in spatially localizing static emitters, tracking multiple emitters over time (usually labeled proteins, or nucleic acids) with comparable resolution to that achieved in widefield superresolution microscopy (SRM) remains an open challenge [4, 5]. Concretely, while photophysical dynamics is leveraged as an asset to widefield SRM in successively localizing nearby emitters in time, it is precisely a disadvantage in tracking as it naturally results in emitters disappearing over several frames. Furthermore, longer data acquisition times per frame (*i*.*e*., longer camera exposures), which otherwise improve the resolution of static emitters within frames, reduce temporal resolution and introduce blurring artefacts for moving emitters [4, 17]. Conversely, short exposures, while necessary in single-particle tracking (SPT), amplify shot noise.

As a compromise allowing for rapid tracking while mitigating the uncertainty introduced by excessive shot noise, it is common to track only a few emitters (or often just one) within a volume roughly the size of a typical bacterium *e*.*g*., Ref. [18], which remains true even when combining photodynamics with tracking in tools such as sptPALM [19]. This is particularly important as small emitters (such as most fluorescent proteins roughly 2 nm in size [20]) result in a diffraction pattern of their visible emitted light over a large region, the point spread function (PSF), within which emitter photons are detected; see Fig. 1a. Given typical conditions under which emitters are tracked, the PSF width, as dictated by the numerical aperture (NA) and the emission wavelength, is about two orders of magnitude larger than the emitter itself, *≈*250 nm. As a result of the PSF’s breadth, when tracking multiple emitters at once in widefield applications, PSFs invariably overlap, obscuring the number and positions of the underlying emitters; see Figs. 1a and 1b. Indeed, widefield SRM was precisely devised to tackle this challenge albeit for static targets by stochastically (PALM/STORM [1–3]) inducing label photophysical transitions to momentarily generate contrast between emitters and their immediate background.

**Figure 1:**
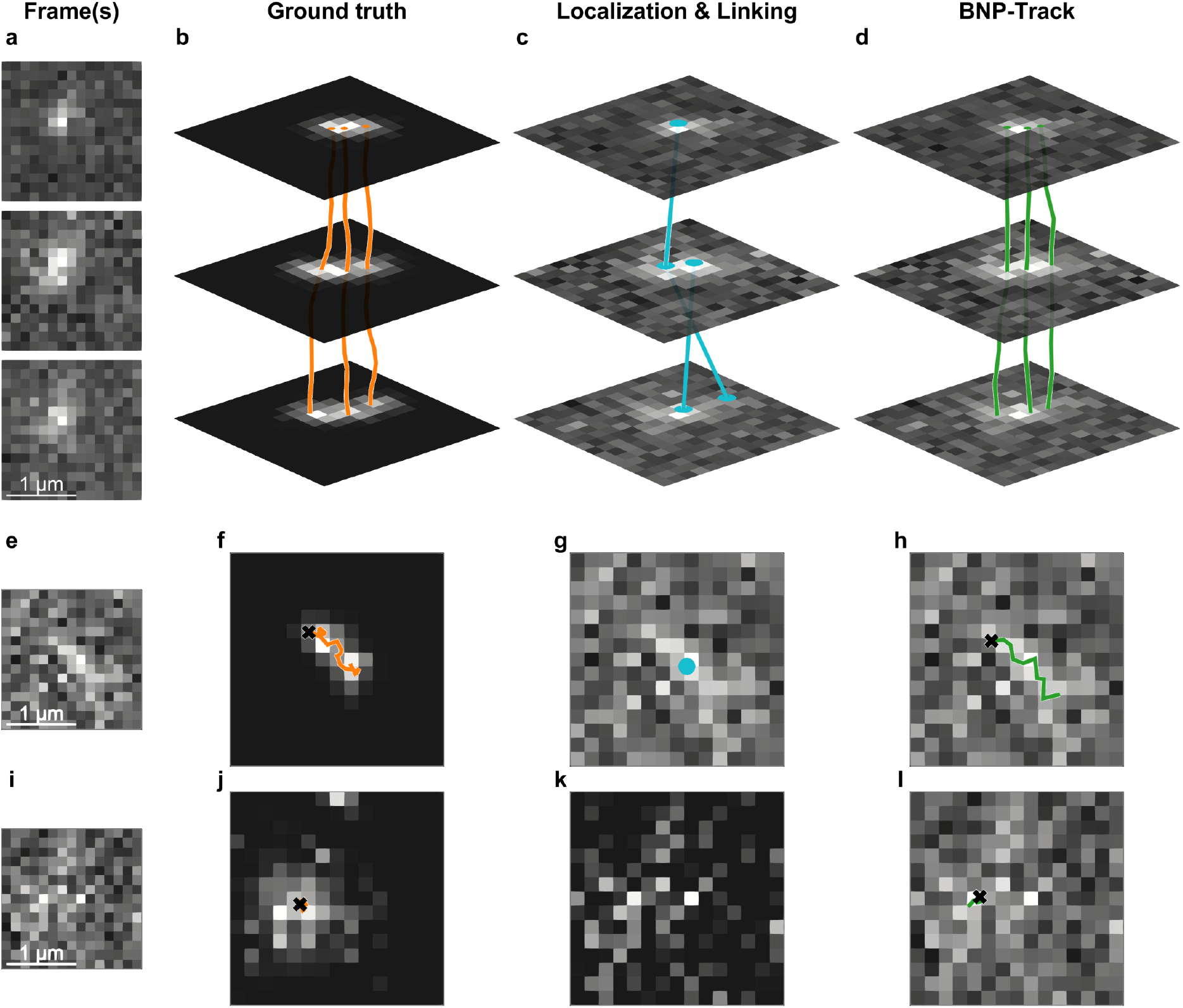
**a**, Three consecutive frames from data generated synthetically following the description provided in Section 3 involving three closely located emitters with the data generated using pixel size 133 nm, NA 1.45, exposure 0.03 s, emission rate 10^4^ s^*−*1^, background flux 10^5^ *µ*m^*−*2^s^*−*1^ and a EMCCD camera; further details provided in Table S.1. **b**, Ground truth tracks shown here in orange superposed to frames where we have removed background for visual appeal. PSF overlap is clear from the three selected frames shown. **c**, Conventional SPT methods often treat the determination of the number of emitters (spot detection) and the ensuing localization and linking of spots as completely or partially separate steps. **d**, BNP-Track leverages all encoded spatiotemporal information to directly infer the number of emitters and their associated tracks across all frames. **e**, A frame of synthetic data with background and a fast-diffusing emitter, all parameters are the same as before except that the diffusion coefficient has now been raised to 1 *µ*m^2^s^*−*1^. **f**, This emitter’s ground truth track (orange) over the course of one frame’s exposure period of duration 0.03 s plotted on top of the noiseless frame, highlighting a highly blurred or “aliased” PSF. The black cross marks the emitter’s location at the end of the exposure. **g**, Conventional localization automatically fails as one single emitter location is not sufficient to approximate a track segment. **h**, BNP-Track considers the emitters’ motion within one exposure (by using information from the present, prior, as well as future exposures) and thereby infers much better tracks. **i**, A frame of synthetic data with an out-of-focus emitter and background. **j**, Similar plot to (**f**). **k**, The frame in (**i**) after performing the difference of Gaussians algorithm [21], a popular prefilter for SPT tools, which in this case removes signal from the emitter, leading to the failure of detecting the only emitter in both (**i**) and (**k**). **l**, By treating the stochastic background within a stochastic framework, BNP-Track manages to track the out-of-focus emitter.

In a companion manuscript we described a tracking tool, BNP-Track, incorporating these features and use it to analyze both *in vivo* and synthetic data motivated from the *in vivo* data sets analyzed. In this manuscript, we focus on four inter-linked aspects of BNP-Track. The first three are explored synergistically while the fourth is discussed in the Methods section Section 4.1.

First, we explore unique features (highlighted in Fig. 1) such as the treatment of intraframe motion over the course of a camera exposure making it possible to extend BNP-Track to longer exposures and faster diffusing emitters. As another example, we also explore BNP-Track’s stochastic treatment of background critical in beating the diffraction limit by capturing the presence of out-of-focus emitters.

Second, related to the latter point, we provide robustness analyses for BNP-Track over parameter regimes (such as variable number of emitters per unit area, diffusion coefficient and photon emission rate) that go beyond the *in vivo* experiments previously explored. We demonstrate on synthetic data (for which ground truth tracks are known) that over reasonable parameter regimes, BNP-Track tracks diffraction-limited emitters with resolution approaching or matching that of SRM for static emitters. Importantly, a recurring theme of our analyses is that a moving emitter distributes its photons in a correlated fashion across both pixels and frames. Thus, information gathered multiple frames ago or even future frames, say, is useful in localizing an emitter in the present frame. Indeed, in our robustness analyses, we even encounter counter-intuitive scenarios where more emitters are sometimes localized to higher resolution than fewer emitters within a field of view (FOV) even when emitter distances fall below the diffraction limit. This is because more emitters may help sharpen our overall estimate for emitter diffusion coefficients.

Third, in the spirit of Ref. [22], we show how BNP-Track’s simultaneous assimilation of the three modular steps of tracking of the existing tracking paradigm (determining emitter numbers, localizing emitters, and linking emitters across frames) allows BNP-Track to outcompete contest-leading localization and linking methods [23–26]. In doing so, we provide approximate head-to-head comparison as exact comparison with other tracking tools is not possible since no method currently simultaneously reports full distributions over emitter numbers and their associated tracks as BNP-Track does. The synthetic data we use in these comparisons includes all effects previously included in Ref. [22] that we also use in the second point to test BNP-Track’s robustness. Naturally, the reason for being selective in our comparison to other tracking tools is also clear. Tracking is a mature field [9, 22–73], with many of the select tools cited reviewed here [5, 22, 74, 75] including by ourselves [4]. Despite employing multiple creative insights balancing computational burden with tracking resolution, we demonstrate through numerical examples in our comparisons that no existing tool can reliably track (below the diffraction limit averaged over the track) more than one protein in a small volume, such as the size of a bacterium’s cytoplasm.

Fourth and finally, we explore the fundamental reason why the existing tracking paradigm cannot beat the diffraction limit. For example, we mathematically demonstrate at a higher level how BNP-Track generalizes statistically-grounded (likelihood-based) tracking methods [64, 66] that require as input the number of emitters (often done in the form of setting localization thresholds). Indeed, we discuss how the more limited assumptions of BNP-Track allow us to extract emitter number which we know already to be encoded in the spatiotemporal correlations across frames currently compromised by the modular, three-step, tracking paradigm.

## 2 Results

Here, we present results on synthetic frame stacks (videos). Unless specified otherwise, the FOV of these videos is about 2 *µ*m by 3 *µ*m for illustrative purposes as we move to denser tracking, mimicking the size of a typical bacterium. Key parameters used in generating the data are listed in Table S.1. Synthetic data generation follows the procedure set by Ref. [22]. For concreteness, the numerical values listed in Table S.1 are motivated by those used in experiments (see citations therein) with widefield illumination using an electron multiplying charge-coupled device (EMCCD) camera. BNP-Track can be readily adapted to any camera by appropriate modification of the emission distribution later detailed in Eq. (8) and any PSF shape by modification of Eq. (6).

After defining the comparison criteria just below, we begin the actual analysis with the simplest scenario, a single emitter, in order to demonstrate that BNP-Track can track one emitter with the same resolution in each frame as existing SRM for static emitters. In this exercise, we use BNP-Track to simultaneously estimate parameters such as the number of emitters (invariably found to be one), diffusion coefficient, background flux, and camera gain. We then analyze a dataset involving three emitters demonstrating that our method can simultaneously track all, even as these move closer to each other than the diffraction limit. The results from these two datasets are then compared with those obtained using TrackMate [72] for performance comparison as TrackMate combines contest-leading single-particle localization and linking methods of Ref. [22].

In the last two sections, we demonstrate how BNP-Track performs over parameter ranges listed in the fifth column of Table S.1. All videos analyzed herein are included in the Supplementary Material.

As a final point, within the Bayesian framework, we do not just report point estimates, but instead present full posterior probability distributions over all relevant quantities, including the number of emitters, spatial tracks (both in and out of focus, to the degree allowed by the PSF’s shape along the axial dimension), the diffusion coefficient, the background flux, and the camera’s gain. To facilitate visual comparison, we also provide *maximum a posteriori* (MAP) estimates for tracks and 95% credible intervals (CIs) for other learned parameters.

### 2.1 Comparison criteria

To better understand performance differences between tracking methods, we provide quantitative metrics using the Tracking Performance Measures tool detailed in Ref. [22]. Here, we provide a brief overview for conciseness. The main measure we report is the pairing distance (per frame per emitter), a metric based on the total gated Euclidean distance between paired tracks averaged over frame number and the ground truth emitter number. When two tracks are further apart than the gate (also referred to as penalty) value *ϵ* at any frame, or a track has missing segments, the distance at that frame is set to be *ϵ*. In the context of this study, *ϵ* is set to be five pixels or 665 nm, which is roughly twice the Rayleigh diffraction limit (0.61*λ/*NA) 280 nm. As there are multiple ways of pairing when comparing two sets of tracks, the final total pairing distance is defined as the minimum pairing distance among all possible pairings of tracks.

It is important to note that the pairing distance metric penalizes linking errors, as it pairs whole tracks during the calculation. Therefore, it should not be directly used to compare with the diffraction limit, which is defined solely based on distinguishing point objects without considering linking across frames. Additionally, as explained in the previous paragraph, the pairing distance also depends on the gate value *ϵ*, which is arbitrarily set by users. When comparing a method’s performance to the diffraction limit, a quantity entirely governed by physics, the performance should not depend on an arbitrary parameter. Thus, the pairing distance is not a suitable metric for comparing to the diffraction limit. Instead, we define the localization resolution, or simply resolution, as the mean Euclidean distance between paired emitter positions across all frames and all emitters. This resolution is equivalent to the pairing distance only if all emitter positions can be paired without any linking errors.

### 2.2 One emitter

The synthetic video we analyzed in this section, Supplementary Video 1, only has one emitter present with results displayed in Fig. 2. To be clear, while we generated the data using one emitter, we still ran BNP-Track which learned that only one emitter was warranted here with negligible probability ascribed to more emitters being present.

**Figure 2:**
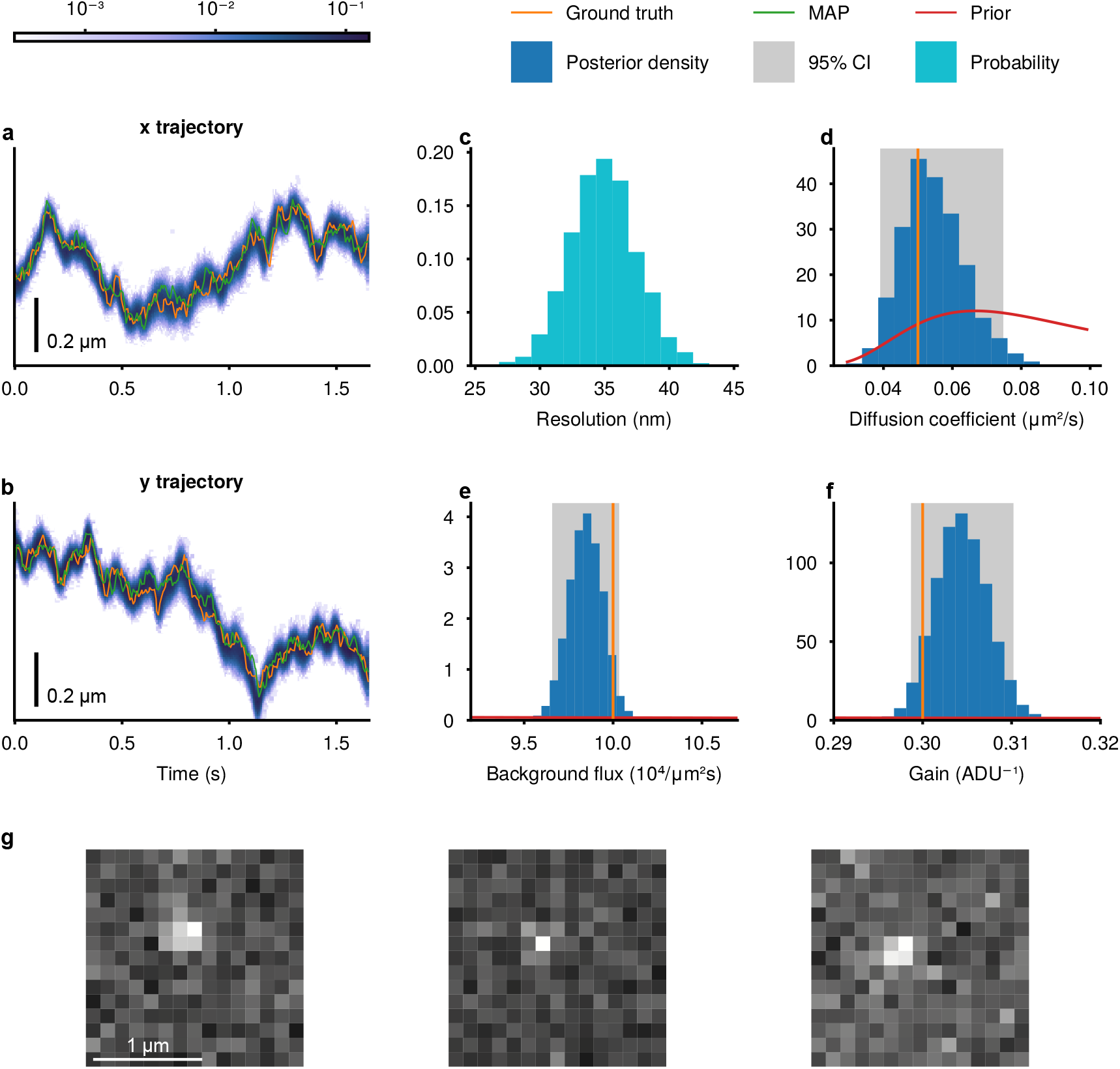
**a** and **b**, BNP-Track’s performance demonstrated using Supplementary Video 1 in terms of the *x* and *y* coordinates of the ground truth tracks compared to BNP-Track’s outputs. The MAP tracks are included for sake of direct comparison. CIs are depicted as shaded regions (which we refer to as CI bands), with darker shading indicating a higher level of confidence. In addition, **c**, The distribution of BNP-Track’s localization resolution distribution (bin heights normalized as probabilities) over all the sampled tracks. See the beginning of Section 2.1 for the definition of localization resolution. **d**-**f**, BNP-Track’s estimate for the diffusion coefficient, background flux, and camera gain (in the case of an EMCCD camera), respectively. The corresponding ground truths and prior probability distributions are also provided. **g**, Three consecutive frames of a selected region from Supplementary Video 1.

Figs. 2a and 2b shows MAP track estimates, ground truth tracks, and the distribution of track samples in *x* and *y*, respectively (see Fig. S.1a for the same plot but for *z*). The darker shaded regions indicate higher confidence and their sizes represent the widths of CI. It is apparent from Fig. 2a that BNP-Track consistently returns a single continuous shaded region surrounding the ground truth track, indicating that BNP-Track correctly identified the emitter number. Additionally, most of the track samples are within *±*100 nm of the ground truth position in each frame. Figure 2b provides a summary metric of the localization resolution. Among all track samples, 95% of the localization resolutions fall between 30.0 nm and 40.0 nm per emitter per frame. This resolution is close to those of the SR localization methods (around 20.0 nm) for static emitters having the same NA, 1.45, as the values of parameters used in simulation, except that we typically use far fewer photons (with only about 250 photons per pixel per frame) [4–10]. Further numerical comparisons with conventional SPT tools are provided in Section 2.4.

In addition to determining tracks, BNP-Track also has the ability to estimate various other parameters, such as the diffusion coefficient, background flux, and camera gain, assuming a EMCCD camera model (though the type of camera model can be generalized). This is demonstrated in Figs. 2d to 2f. The ground truth values for each of these parameters fall within their corresponding 95% confidence intervals. Specifically, the ground truth values and confidence intervals are as follows: 0.05 *µ*m^2^s^*−*1^ vs. (0.039 to 0.075) *µ*m^2^s^*−*1^ for the diffusion coefficient; 10^5^ *µ*m^*−*2^s^*−*1^ vs. (0.966 to 1.003) *×*10^5^*µ*m^*−*2^s^*−*1^ for background flux; 0.3 ADU^*−*1^ vs. (0.299 to 0.310) ADU^*−*1^ for the camera’s gain.

### 2.3 Three emitters

As highlighted earlier, the ability to handle multiple emitters within the FOV is crucial in tracking. Figure 3 demonstrates BNP-Track’s performance sharing all the same parameters as the single emitter case in Fig. 2 except for three emitters. The synthetic data is provided in Supplementary Video 2. As shown in Fig. 3a, BNP-Track accurately learns the number of emitters by returning track samples distributed as CI bands that we find to be centered at the ground truth even as these move in and out of focus; see Fig. S.1b for the same plot but for *z*.

**Figure 3:**
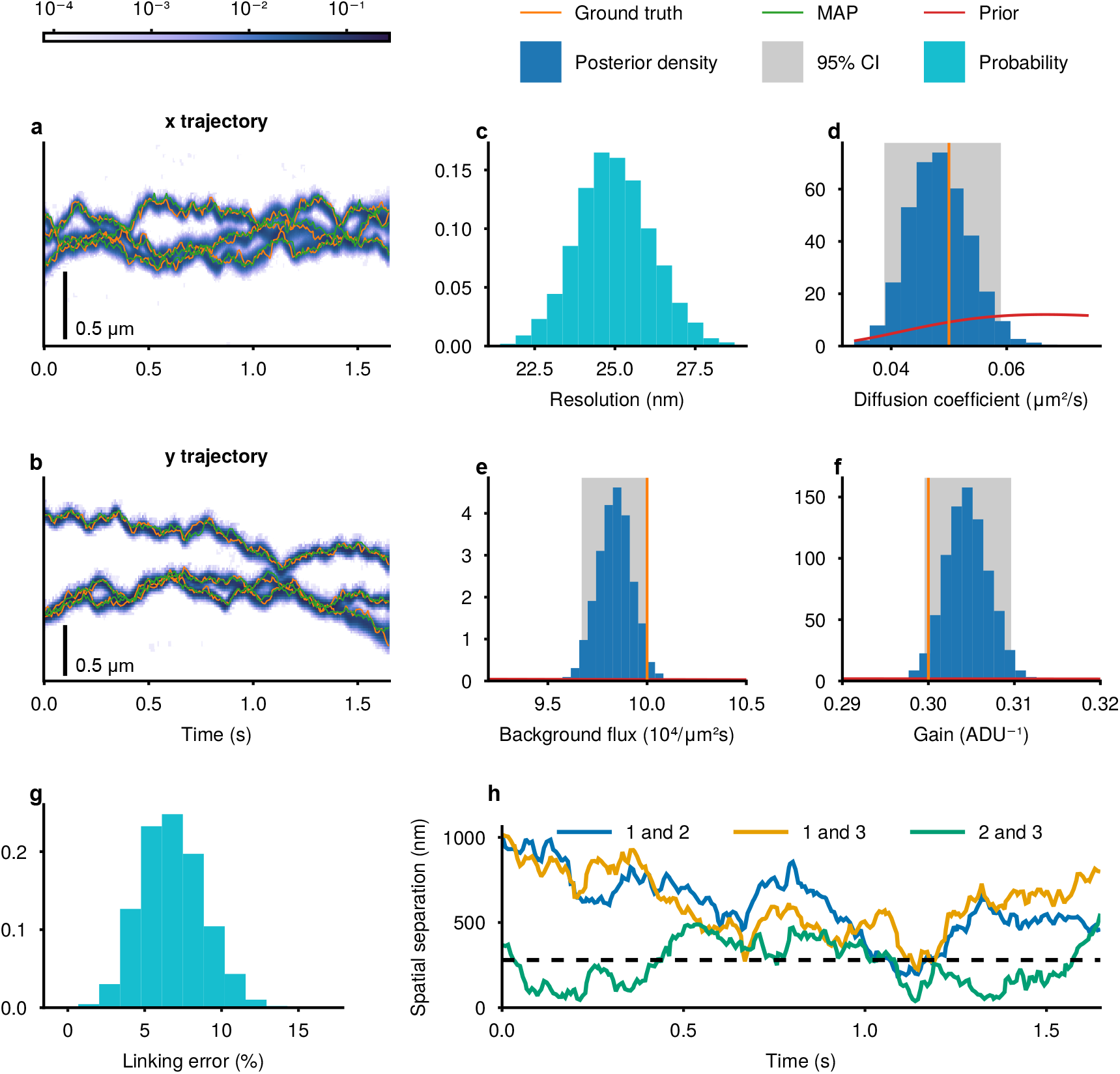
BNP-Track’s performance demonstrated using a synthetic dataset with three emitters (Supplementary Video 2). The layout is similar to Fig. 2 but with two additional panels: **g**, The distribution of incorrect linking percentages; **h**, The spatial separation between each emitter pair as a function of time and the Rayleigh diffraction limit, is marked by the black dashed line.

One unique challenge in tracking multiple emitters is the occurrence of criss-crossing tracks, where the separation between emitters approaches or falls below their typical PSF widths. For example, this is the case in Supplementary Video 2, where two out of the three emitters move closer to each other than the diffraction limit; see Fig. 3h. As we will explore shortly, these scenarios are especially challenging for the existing three-step tracking paradigm.

As a benchmark for tracking multiple emitters, we counted the number of incorrect links (across frames) over the entire video for all sampled tracks; the results are shown in Fig. 3g. Here 95% of all tracks sampled from our joint posterior over emitter numbers and tracks had between approximately 2.7% and 10.9% incorrect links; see Fig. S.2 for another three-emitter video (Supplementary Video 3) but with emitters even closer to each other in space.

Similar to the one-emitter case, we also calculated the 95% CI for the localization resolution, which was found to be 22.7 nm to 27.3 nm, and thus on par with SRM for static emitters under similar exposure, emission rate, background, and gain (see Fig. 3c). Perhaps initially counter-intuitive, we note that this range is actually lower than the localization resolution CI calculated in the one-emitter video (Supplementary Video 1), 30.0 nm to 40.0 nm. Indeed, one might expect higher uncertainty when more emitters are present in the FOV. In general more precise localization in the presence of more emitters is possible, fortuitously, if these emitters happen to spend more time on the focal plane where they are localized more accurately. This is not the case here; we verified that the average distances between emitters and the focal plane in Video 1 and 2 are not substantially different. Rather, we found that with more tracks, we have more spatiotemporal correlation that BNP-Track leverages to refine the estimate of some parameters, say diffusion coefficients. This hypothesis is supported by the fact that BNP-Track’s output in the three-emitter video (Fig. 3d) shows a narrower 95% CI for the diffusion coefficient, (0.039 to 0.059) *µ*m^2^s^*−*1^, compared to the one-emitter video, (0.039 to 0.075) *µ*m^2^s^*−*1^.

While gathering more spatiotemporal correlation from tracks can help us infer emitter tracks (by improving diffusion coefficient estimates), estimating (homogeneous) background flux and gain remains unaffected as they are assumed time-independent here. Indeed, consistent with this expectation, in Figs. 3d and 3e we present BNP-Track’s estimates of these two parameters: (0.967 to 1.001) *×* 10^5^*µ*m^*−*2^s^*−*1^ for background flux’s 95% CI; (0.300 to 0.310) ADU^*−*1^ for camera’s gain’s 95% CI.

It is worth noting here that BNP-Track actually explored the possibilities of assigning different number of emitters. This can not be seen directly from the number of CI bands in Fig. 3 as the corresponding chances remain almost negligible, see Fig. S.3h which involves Supplementary Videos 1, 2, and 4.

### 2.4 Comparison with TrackMate

It is difficult to directly compare BNP-Track to other SPT methods head-to-head as no existing method currently estimates (full distributions over) both emitter number, alongside their associated tracks. Thus, we necessarily must provide an advantage to existing methods by providing them with correct emitter numbers ahead of time. As we will see, even under this optimistic scenario, BNP-Track still exceeds the resolution of existing tools and yields reduced error rates (percentage of wrong links).

With this *proviso*, we now compare the tracking performance of BNP-Track and TrackMate [72], the standard SPT tool which houses several detection and tracking modules tested in Ref. [22], by analyzing the same datasets and subsequently comparing the inferred tracks against the ground truth using the Tracking Performance Measures tool [22] in Icy [76].

To prepare the benchmark data, we used both one- and three-emitter videos from Supplementary Videos 1 and 2. In order to compare both methods, as BNP-Track provides a full posterior over tracks and emitter numbers while TrackMate provides only best track estimates (according to criteria we discuss shortly), we selected as point of comparison BNP-Track’s MAP tracks. The latter tracks maximize the full joint posterior probability distribution and can therefore be considered the overall “best” tracks.

On the other hand, what is chosen as the best track according to conventional SPT tools depends on the application at hand. This arises for two main reasons: 1) without a numerical criterion like a posterior, decisions as to which track is better depends on pre-selected metrics (*e*.*g*., tracks with minimal spurious detections or tracks with the fewest missed links); 2) although it is generally desirable to have tracks with no false positives (spurious detections or tracks) and no false negatives (missed detections or tracks), it is often difficult to optimize both, as most of these methods do not allow frame-specific thresholds. Therefore, when false negatives and false positives cannot be reduced to zero simultaneously, we use TrackMate to generate two sets of single-particle tracks: Set A which prioritizes minimizing false positives in the localization (spot detection) phase, followed by minimizing false negatives in the linking phase, while Set B prioritizes minimizing false negatives in the localization phase, followed by minimizing the other type of errors in the linking phase.

For more information on the TrackMate specific input parameters (such as PSF width estimate and gap closing distance), see Section 4.4. We also note that, in order to maximally challenge BNP-Track, we provided strong advantages to TrackMate by performing both aforementioned error minimization steps using ground truth emitter number in the localization step and ground truth diffusion coefficient in the linking step.

Fig. 4 provides a direct visual comparison between the ground truth, BNP-Track MAP track and TrackMate tracks for the benchmark videos. In the one-emitter case, both BNP-Track (Fig. 4a) and TrackMate (Fig. 4b) successfully track the single emitter throughout the entire video with similar resolution (34.6 nm vs. 35.4 nm). In order to further demonstrate that conventional SPT tools would typically work for this relatively simple scenario, we also include the track obtained with u-track [25] using the point source particle detection process (Fig. 4c), which again, achieves a similar resolution at 32.1 nm. We note that in Figs. 4a to 4c, resolution is equal to pairing distance as there is no missing segments or incorrect links in tracks.

**Figure 4:**
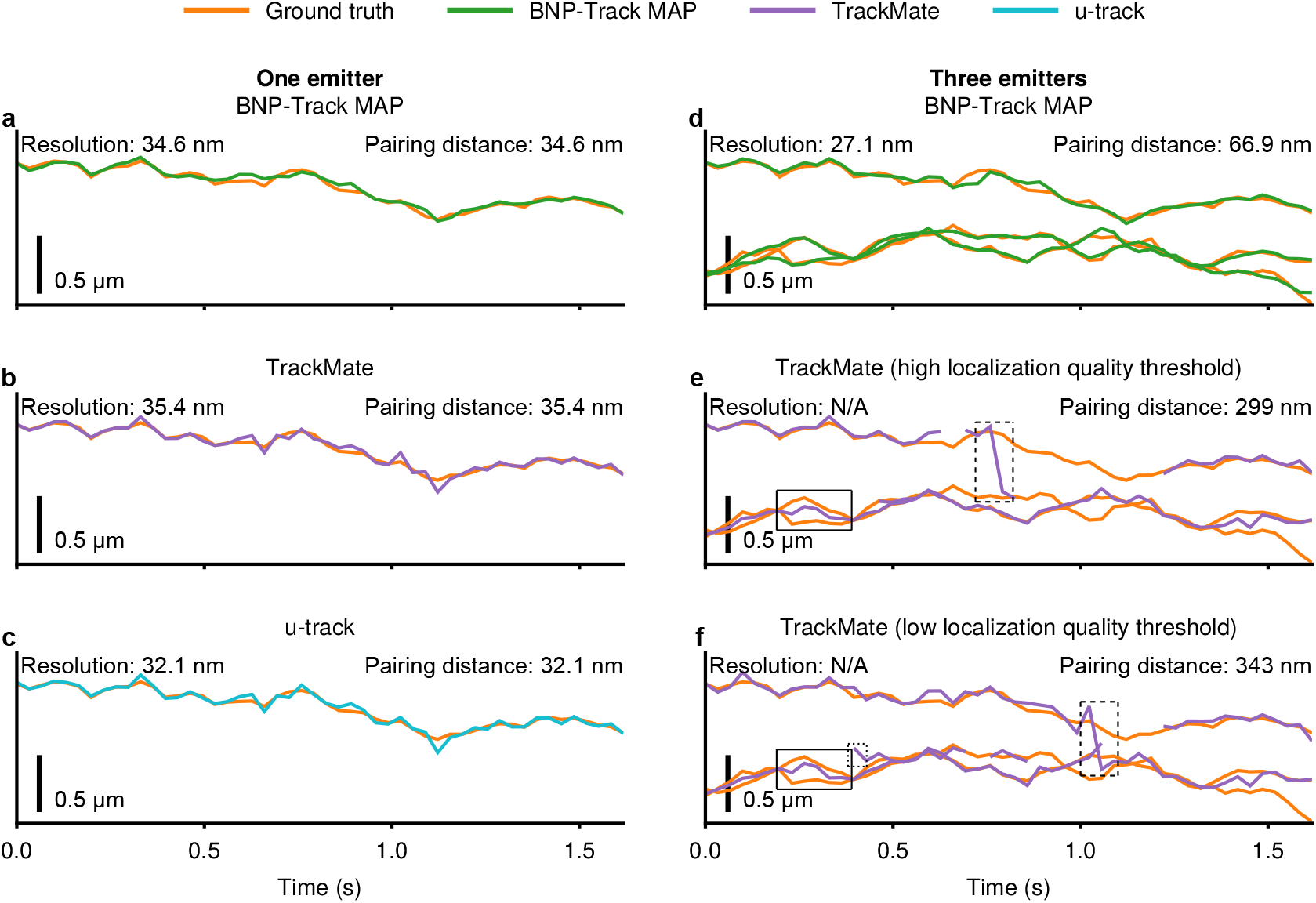
A comparison of tracking performance among BNP-Track, TrackMate, and u-track using two synthetic datasets with one emitter (Supplementary Videos 1) and three emitters (Supplementary Videos 2) in the *y* coordinate. See Fig. S.4 for the same figure but with the *x* coordinate. **a**, BNP-Track’s MAP estimates compared with the one-emitter ground truth. **b**, TrackMate’s estimates compared with the one-emitter ground truth. **c**, u-track’s estimates compared with the one-emitter ground truth. **d**, BNP-Track’s MAP estimates compared with the three-emitter ground truth. **e** and **f**, TrackMate’s estimates compared with the three-emitter ground truth with high and low localization quality thresholds, respectively. In (**d**-**f**), the top ground truth track is the same as the ground truth in (**a**-**c**). The boxed regions in (**e**) and (**f**) highlight where TrackMate performs relatively poorly.

Predictably, as we begin encountering PSF overlap in multiple emitters for the three-emitter dataset, BNP-Track and TrackMate’s performance diverge (we note that u-track’s performance is not shown here as the current FOV is too small for running its algorithm). While BNP-Track remains capable of tracking all three emitters throughout the entire video with a resolution of 27.1 nm from the MAP tracks, even as these move below diffraction limit in frames 2 to 13 and 34 to 47 (see Fig. 3h), several issues arise for the TrackMate tracks (Figs. 4e and 4f). For instance, diffraction limited emitters get interpreted as one emitter (solid boxes), incorrect links with large jumps (dashed boxes), and spurious detections (the dotted box) from TrackMate set B. These issues indicate that TrackMate can no longer resolve the emitters in this dataset, and hence no resolution is calculated. Therefore, for the sake of a quantitative comparison, we calculated the pairing distance, defined in Section 2.1, between the ground truth tracks and each SPT method’s output: BNP-Track’s MAP tracks have a pairing distance of 66.9 nm while both TrackMate track sets yield pairing distances no less than 300 nm.

Two additional points should be noted here: 1) as illustrated in Fig. 4d, BNP-Track’s pairing distance is larger than its resolution, which is due to linking error. Nevertheless, we argue this is not a major concern since it only occurs near frame 35 where two emitters are less than 40 nm apart (Fig. 3h), and hence, potential further analyses (such as learning diffusion dynamics) are not affected; 2) Even though the three-emitter dataset contains the same track as the one-emitter dataset, BNP-Track actually achieves a better resolution in the more complicated three-emitter dataset. We attribute this to the fact that BNP-Track leverages all spatiotemporal correlation detailed in the next section.

### 2.5 Benefits of leveraging all spatiotemporal correlation

To showcase the advantages of utilizing all spatiotemporal correlation, we generated another one-emitter dataset intentionally consisting of a large number of frames (200) for illustrative purposes, which can be viewed in Supplementary Video 5. We initially applied BNP-Track to only the first two frames and then progressively incorporated more frames into the analysis. We monitored BNP-Track’s performance at each iteration for the arbitrarily chosen frame 2, and this procedure can be repeated for any other frame of interest.

BNP-Track’s resolution at frame 2 improves with the inclusion of more future frames in the analysis, as depicted in Fig. 5b. This is in contrast to any SPT method that separates localization and linking as modular steps [9, 23–53, 72]. The outcome suggests that BNP-Track can obtain spatiotemporal correlation from “future” (and “past”) frames, thereby enhancing the resolution at the “current” frame. The additional information is transmitted through the system’s diffusion dynamics, also better assessed by taking more frames into account, as shown in Fig. 5c. Including more frames is not the only way to increase the amount of information contained in a dataset: the presence of more emitters has the same effect. This explains why BNP-Track’s resolution improves from Fig. 4a to Fig. 4d. In other words, frames and numbers of emitters all provide information and, as such, similar information may be gathered by considering more emitters even in the presence of far fewer frames than 200 (as would be typical for most tracking experiments).

**Figure 5:**
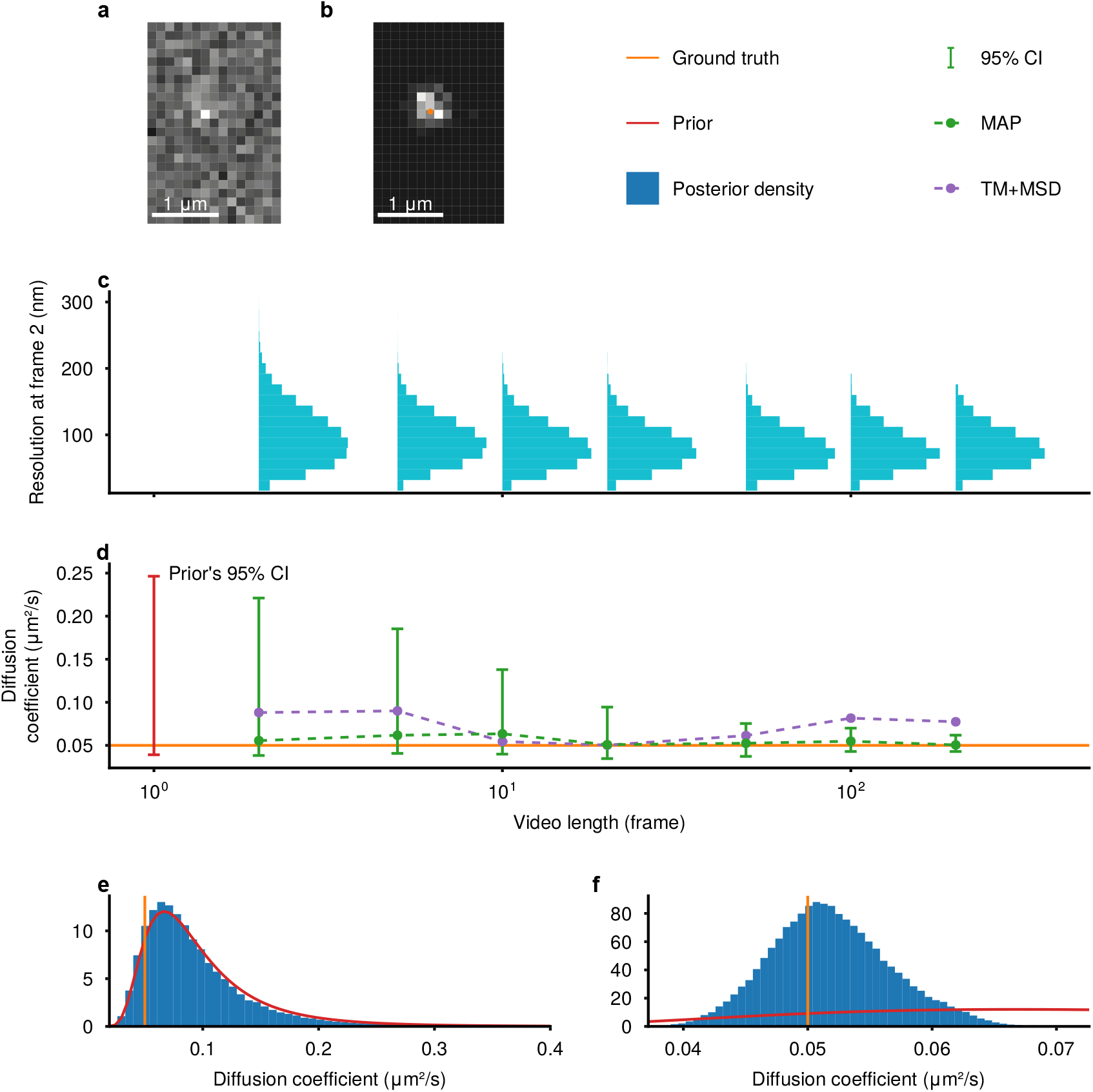
BNP-Track’s performance as the length of the video increases. **a**, The second frame of Supplementary Video 5. **b**, The same frame without noise, with the ground truth position of the emitter marked as the orange dot. **c**, The localization resolution in this frame versus the number of frames considered in the analysis. **d**, The diffusion coefficient’s MAPs and 95% CIs as a function of the frame number used in the analysis, compared with the values inferred by TrackMate and mean squared displacement (TM+MSD in the legend). The diffusion coefficient estimate also improves as more frames are analyzed. Note that, as TrackMate was manually tuned to obtain the best track, the performance of TM+MSD in (**d**) is artificially improved. Even in this case, BNP-Track still produces more accurate and consistent estimates. **e**, The diffusion coefficient’s posterior probability distribution inferred by BNP-Track with only the first two frames, indicating a lack of information as the prior and posterior are similar. **f**, The diffusion coefficient’s posterior probability distribution inferred by BNP-Track with all 200 frames in Supplementary Video 5.

Leveraging all spatiotemporal correlations also results in more efficient information utilization. This is demonstrated in Fig. 4c, where it can be observed that BNP-Track’s estimate of the diffusion coefficient converges more rapidly with respect to the number of frames analyzed than that obtained using TrackMate combined with mean squared displacement.

### 2.6 Robustness tests

In this section, we evaluate the robustness of BNP-Track under different parameter ranges, including emitter numbers, diffusion coefficients, (photon) emission rates, and background flux. That is, we start with default parameter values set in the fourth column of Table S.1 motivated from literature values and vary one parameter at a time over the ranges listed in the last column of the same table. We note that the tests with the standard parameter values are already illustrated in Figs. 2 and 3.

We first test BNP-Track’s ability to handle multiple emitters by introducing seven emitters within the same FOV (2 *µ*m *×* 3 *µ*m) as before (Supplementary Video 4); see Fig. S.3. Despite this dramatic increase in complexity, BNP-Track assigns over 94% probability to seven emitters with a pairing distance of 55.5 nm, much smaller than TrackMate’s 323 nm.

BNP-Track is also robust when tested on videos generated with different diffusion coefficients, as shown in Fig. 6. Here, BNP-Track can accurately track all emitters, as determined by localization resolution discussed shortly, and determine the correct diffusion coefficient, even when the diffusion coefficient is five times higher or lower than the reference value. As observed in Fig. 6, the distribution of emitter positions becomes broader as the diffusion coefficient increases. This is supported by the 95% confidence intervals of the localization resolutions, which range from (15.2 to 18.8) nm in the first row, (22.8 to 27.3) nm in the second row, (47.8 to 65.4) nm in the third row. One major factor contributing to this trend is motion blur introduced by fast diffusion coefficients. Another important factor is that faster diffusing species have less time to remain within the field view or move away from the in focus plane, resulting in fewer informative frames (frames with more detected photons). To further understand the impact of motion blur on BNP-Track’s results, we also conducted further testing illustrated in Fig. S.5.

**Figure 6:**
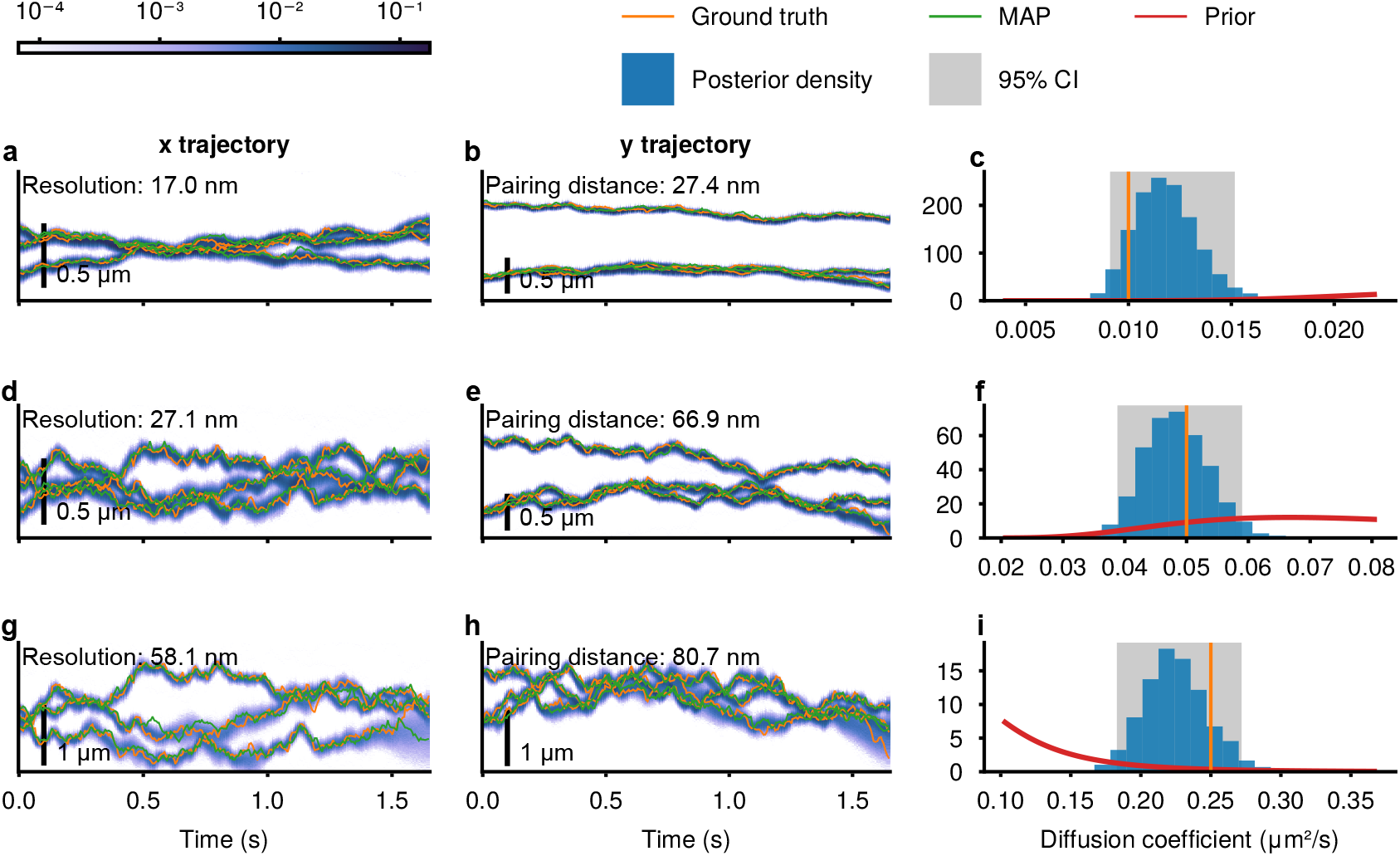
BNP-Track’s performance under various diffusion coefficients. **a**-**c**, BNP-Track’s analysis for Supplementary Video 6 with 0.01 *µ*m^2^s^*−*1^. **d**-**f**, BNP-Track’s analysis for Supplementary Video 2 with 0.05 *µ*m^2^s^*−*1^. **g**-**i**, BNP-Track’s analysis for are from Supplementary Video 7 with 0.25 *µ*m^2^s^*−*1^.

We performed similar robustness tests by varying the photon emission rate of the emitters and background photon flux. These two parameters both affect the signal-to-noise ratio and, as such, we present their results together. As shown in Figs. S.6 and S.7, BNP-Track successfully tracks all three emitters. As expected, low emission rates and high background flux worsen both resolution and pairing distance. Once again, the localization resolution 95% CIs are provided as an overview: (22.7 to 27.3) nm for emission rate 10^4^ s^*−*1^, background flux 10^5^ *µ*m^*−*2^s^*−*1^; (54.7 to 76.7) nm for emission rate 0.2 *×* 10^4^ s^*−*1^, background flux 10^5^ *µ*m^*−*2^s^*−*1^; (8.5 to 10.3) nm for emission rate 5 *×* 10^4^ s^*−*1^, background flux 10^5^ *µ*m^*−*2^s^*−*1^; (16.8 to 19.9) nm for emission rate 10^4^ s^*−*1^, background flux 0.2 *×* 10^5^ *µ*m^*−*2^s^*−*1^; (34.6 to 43.6) nm for emission rate 10^4^ s^*−*1^, background flux 5 *×* 10^5^ *µ*m^*−*2^s^*−*1^.

Additionally, in order to demonstrate the necessity of using BNP-Track’s feature to infer emitter numbers, we also performed two tests where the emitter number was fixed albeit to incorrect values. See Figs. S.9 and S.10 for how this impacts BNP-Track’s performance. Briefly, forcing BNP-Track to run with an emitter number below that set by ground truth naturally causes jumps between tracks, resulting in over estimating diffusion coefficient by a factor of ten. On the other hand, running with more emitters typically is less problematic as BNP-Track will treat the extra emitters as out of the FOV.

More than its robustness across various parameter ranges, it is worth examining whether BNP-Track can be applied to systems with different motion models, given that it currently only considers the Brownian motion model. We address this concern in our companion manuscript, where we analyze an experimental dataset with an unknown motion model.

## 3 Discussion

Here, we introduced BNP-Track, a tracking tool capable of superresolved tracking. That is, BNP-Track tracks moving emitters with approximately as few as 250 photons per pixel in each frame (with localization resolution comparable to SRM) and tracks individual emitters even as these come closer to one another than light’s diffraction limit.

To achieve this, BNP-Track abandons the existing tracking paradigm which relies on three separate/modular operations: emitter number determination, single-emitter localization, and linking. Rather, BNP-Track leverages all spatiotemporal correlation available in order to simultaneously obtain a full joint posterior over emitter numbers, their associated tracks, diffusion coefficients, background photon flux, and camera gain (assuming EMCCD).

The capability to track as many as ten or more small molecules within an area of about 5 *µ*m^2^ may help provide direct insight on transient co-localization of emitters [77], as well as biomolecular cluster (dis)assembly [78] beyond the reach of existing tracking paradigm.

In addition to its robust tracking resolution over a range of reasonable parameters, BNP-Track’s software suite is tailored for a graduate student end-user in mind. Here users need as their only input calibrated parameters including the objective’s NA, refractive index, emission wavelength, photon emission rate, pixel size, and exposure time. Other tracking tools such as TrackMate require a significant amount of fine-tuning and thresholding.

While BNP-Track has the ability to achieve high tracking performance and simultaneously infer relevant parameters, it does have higher associated computational time. For example, in the three-emitter video discussed in this paper, while TrackMate takes only a few seconds to complete localization and linking, BNP-Track requires approximately a day on a mid-range desktop computer. Additionally, BNP-Track’s time consumption increases roughly linearly with the number of emitters. However, we argue here that this is not a significant drawback for these reasons: First, users must often spend considerable time tuning the system to use TrackMate or similar tools, which exceeds the computation time. More tuning is then required for slightly different imaging conditions. In contrast, BNP-Track requires more computational time (wall time) but less manual effort. Second, for systems with well-calibrated background flux and/or gain, BNP-Track can use these values as input and save the time that would be required to learn these (which would reduce the computational time for the three emitter case by about 30%). Finally, BNP-Track can also take the outputs of other SPT tools as a starting point (*i*.*e*., the initial Monte Carlo seed) and perform analysis on top of it, reducing the computational time required from a day to several hours and sometimes even less.

BNP-Track, in itself, is a first and foremost a mathematical and conceptual framework and the framework highlighted in Section 4 can be generalized to include various camera models (*e*.*g*., sCMOS versus EMCCD) or any pre-calibrated form of the PSF however aberrated. Other generalizations include extending the camera model to accommodate different experimental setups, including multicamera imaging for improved lateral resolution, multiplane imaging for improved axial resolution, or multicolor imaging for Förster resonance energy transfer labels. Similarly, we may incorporate the effects of a vanishing illumination field from total internal reflection fluorescence microscopy or techniques such as HILO microscopy [79].

## 4 Method

### 4.1 Conceptual basis

Before we describe details of BNP-Track, we provide a discussion on the fundamental design reasons why BNP-Track surpasses conventional tracking methods.

To do this, we first establish some basic notation. Consistent with our companion manuscript, the set of emitter positions of all *M* emitters across *N* frames are denoted as 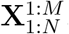, and the measurements of all pixels across all frames are denoted as 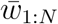. The ultimate goal of single-particle tracking methods is to determine the optimal set of 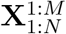 as well as *M* given 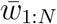 (or, better yet, the full distribution over these quantities given experimental uncertainty). In all tracking methods, an optimization criterion is defined by a function dependent on tracks and measurements, expressed as 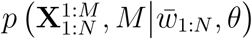, where *θ* includes all relevant parameters such as the emitters’ diffusion coefficient.

In a Bayesian framework, 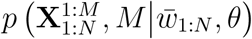 is referred to as the posterior probability distribution and represents “the probability distribution of 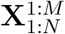 and *M* given 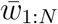 and *θ*”. The set of tracks globally maximizing the posterior is the “best”. In the following paragraphs, we demonstrate how conventional tracking methods, treating localization and linking as separate steps, approximate *p*, while BNP-Track does not. This feature ultimately limits the resolution of existing methods.

We begin by factorizing *p* for each frame. Bayes’ theorem allows for the following factorization without approximation

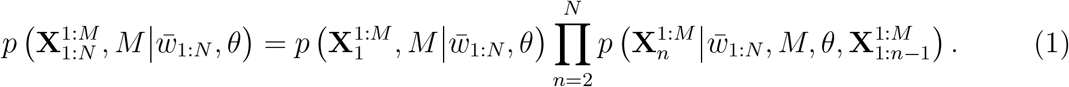

Writing 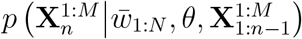 means that localizing emitters on frame *n* in the Bayesian framework takes not only measurements but also emitter positions at every frame from the past into account, as all positions are correlated through the emitters’ motion model. In fact, emitter positions from all future frames contribute in the same way. If this is not immediately clear, *p* can be exactly refactored as

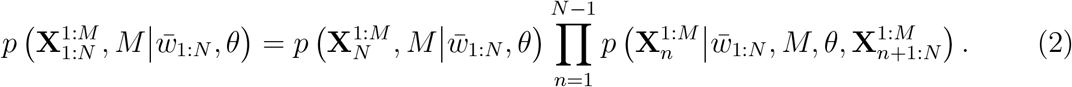

Another common factorization is

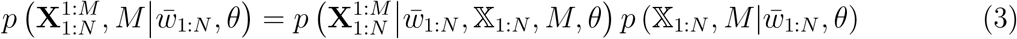

where 𝕩_*n*_ represents all detected emitter positions at frame *n* before they are assigned an emitter label. This means that 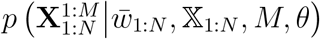 treats linking conditioned on localization while 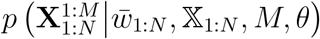 treats localization alone.

The factorization methods described in Eqs. (1) to (3) are equivalent and retain all spatiotemporal information, enabling them to achieve the highest possible tracking resolution if global optimization is performed which, so far, has remained a longstanding challenge. BNP-Track addresses this issue by utilizing novel computational statistics techniques, while conventional tracking methods resort to approximations in order to proceed.

A common approximation used is to separate localization and linking into separate steps [9, 23–53, 72]. Using notation established in Eq. (3), this is equivalent to first identifying those 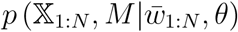 and *M* ^*^ optimizing 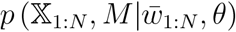, and then optimizing 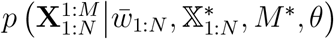 In other words, separating localization and linking within the framework of a greedy algorithm not guaranteed to identify global optima.

Moreover, as the total number of emitters, *M*, is necessarily determined by jointly considering all frames, optimizing 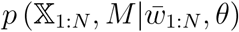 by itself becomes difficult. Therefore, most emitter localization methods [80–82] further approximate

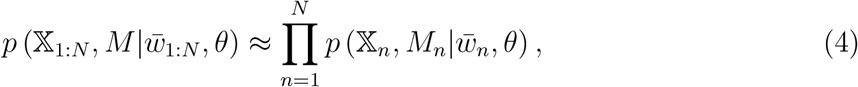

allowing frames to support different emitter numbers. However, in order to incorporate different *M*_*n*_ ‘s, the optimization of 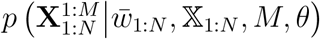 must also be modified accordingly, for example, by allowing gap closing [9, 25, 26, 29, 39–41, 43, 46–48, 50, 53, 64, 72]. All these (generally difficult to control) approximations and modifications, absent in BNP-Track, eventually degrade tracking resolution.

### 4.2 Data synthesis

Briefly summarizing the companion manuscript, we consider *N* frames in each synthetic video and, for each frame, we select equally spaced time points within the exposure period between the previous frame and the current one. For example, for frame *n*, we have time points *{t*_*n*,1_, *t*_*n*,2_, …, *t*_*n,K*_*}*, where *K* is the total number of time points. We also assume that there are *B* (*a priori* unknown) emitters undergoing Brownian motion with constant diffusion coefficient *D*. The spatial position of emitter, 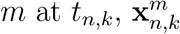, is determined by sampling from the following multivariate Normal distribution denoted as **Normal**_3_

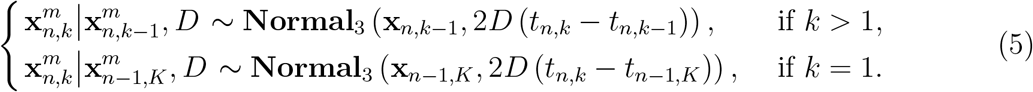

For concreteness here, the contribution of each 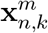 to pixel *p* is calculated using a circular Gaussian Lorentzian PSF [83] though any pre-calibrated form is easily accommodated. The total photon contribution from emitter *m* to pixel *p* within frame *n* is denoted as 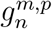 and is obtained by integrating *m*’s PSF over the area and exposure time of pixel *p*

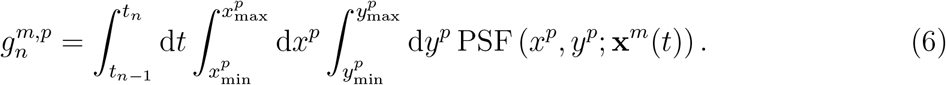

The calculation of this integral is described in Section 3. The total number of photons received by pixel *p* within frame *n*, denoted as 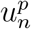, is then given by

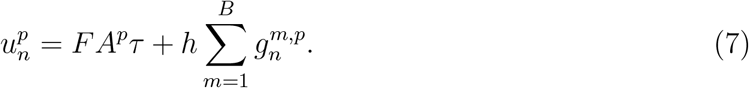

Here, *F* represents the background photon flux, *A*^*p*^ is the area of pixel *p*, and *h* is the photon emission rate. In this study, we consider an electron-multiplying charge-coupled device (EMCCD) camera, whose readout is modeled by

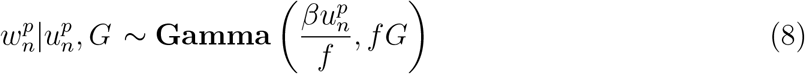

where *f* is the noise excess factor (equals 2 for EMCCDs), *β* is the quantum efficiency, and *G* is the gain. A more detailed description of these parameters can be found in Section 3.

### 4.3 Inference model

As previously mentioned in this manuscript and the companion manuscript, our goal is to estimate the collection of emitter tracks, represented as

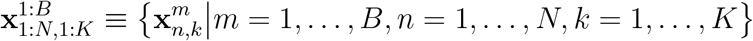

as well as the number of emitters *B*, the diffusion coefficient *D*, the background flux *F*, and the EMCCD gain *G* if necessary, provided all measurements

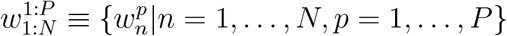

. Within a Bayesian framework, this implies that we wish to sample from the posterior probability distribution 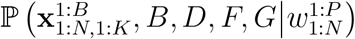.

However, this nonparametric posterior does not assume a simple form and must be simulated computationally. Rather than inferring *B* directly, we consider a total *M* emitters with *M ≫ B*. Here each emitter is labeled with a binary value, *b*^*m*^, known as the load. A value of *b*^*m*^ = 0 indicates that the emitter does not emit any photons (*i*.*e*., the load is inactive or, equivalently, the emitter is unnecessary in explaining the data), while *b*^*m*^ = 1 signifies that the load is active (again, equivalently, that the emitter is warranted by the data). This allows us to compute *B* as the sum of all *b*^*m*^ values. Therefore, the target posterior becomes 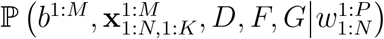

Using Bayes’ theorem, we construct the posterior from the product of the observation likelihood, 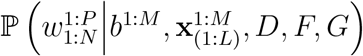, and prior, 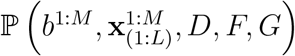. Both terms are detailed in Section D.1. From the posterior constructed from this product, we may start sampling the posterior to deduce appropriate values over the parameters.

In order to sample this high dimensional posterior, we develop a specialized Markov Chain Monte Carlo (MCMC) scheme. In particular, a global Gibbs sampling [84] scheme is combined with Metropolis-Hastings (MH) algorithm [85, 86] and ancestral sampling to sample the posterior. Within the Gibbs sampler of BNP-Track, the parameters are updated in the following order:

1. Update *G* by sampling from its marginal posterior 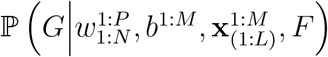 (Section D.2);

2. Update *F* by sampling from 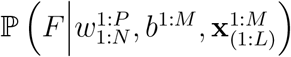 using the MH algorithm (Section D.3);

3. Update *b*^1:*M*^ by sampling from 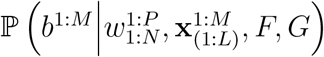 (Section D.4);

4. Update 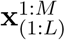 and *D* by sampling from 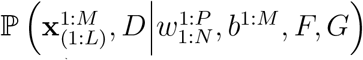 using the MH algorithm and ancestral sampling (Section D.5);

5. Update 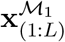 again by sampling from 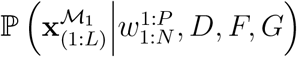 (Section D.6).

These steps are repeated until the sampler has reached convergence.

All algorithms describe above are implemented in MATLAB R2022b. Figures are generated by Makie [87] v0.19.2 using Julia [88] v1.8.5.

### 4.4 TrackMate and tracking performance measures

Besides its widespread use, ongoing maintenance and updates, and being built upon leading methods in Ref. [22], we opted for TrackMate specifically because it offers various localization and linking methods, as well as multiple thresholding options.

To generate tracks for comparison in 2.4, we first exported synthesized videos to TIFF files and imported them in Fiji [89] v1.54b for analysis with TrackMate [72] v7.9.2. As part of implementing TrackMate, the Laplacian of Gaussian detector with sub-pixel localization and the linear assignment problem (LAP) mathematical framework [25] were used in spot detection. Spots were then filtered based on quality, contrast, sum intensity, and radius. For the LAP tracker, we allowed gap closing and tuned the parameters of max distance, max frame gap, and number of spots in tracks to find the best tracks. No extra feature penalties were added.

The benchmarks in Tables S.4 and S.5 were created using the Tracking Performance Measure plugin in Icy 2.4.3.0. To generate these benchmarks, we exported the TrackMate track, the BNP-Track MAP estimates, and ground truth tracks as XML files. All tracks were imported into Icy’s TrackManager using the “Import TrackMate track file” feature, and the Tracking Performance Measure plugin was initiated using the “add Track Processor” option. The only required input for this plugin is the “maximum distance between detections”, which we kept at a default value of five as this value is large enough for the diffusion coefficient in the synthetic data (0.04 *µ*m^2^s^*−*1^).

## 5 Acknowledgements

We thank NIH NIGMS (R01GM130745) for supporting early efforts in nonparametrics and tracking, NIH NIGMS (R01GM134426) for supporting single photon efforts, and NIH MIRA entitled “Toward high spatiotemporal resolution models of single molecules for in vivo applications”. L. X. thanks Mohamadreza Fazel, Ayush Saurabh, Shep Bryan, Maxwell Schweiger, and Alex Rojewski for helpful discussions.

## 6 Supporting Material

This section is attached separately.

## References

1. Betzig, E. et al. Imaging Intracellular Fluorescent Proteins at Nanometer Resolution. Science 313, 1642–1645. eprint: https://www.science.org/doi/pdf/10.1126/science.1127344. https://www.science.org/doi/abs/10.1126/science.1127344 (2006).

2. Rust, M. J., Bates, M. & Zhuang, X. Sub-diffraction-limit imaging by stochastic optical reconstruction microscopy (STORM). Nat. Methods 3, 793–796. issn: 1548-7105. https://doi.org/10.1038/nmeth929 (Oct. 2006).

3. Hess, S., Girirajan, T. & Mason, M. Ultra-High Resolution Imaging by Fluorescence Photoactivation Localization Microscopy. Biophys. J. 91, 4258–4272. issn: 0006-3495. https://www.sciencedirect.com/science/article/pii/S0006349506721403 (2006).

4. Lee, A., Tsekouras, K., Calderon, C., Bustamante, C. & Pressé, S. Unraveling the Thousand Word Picture: An Introduction to Super-Resolution Data Analysis. Chem. Rev. 117. PMID: 28414216, 7276–7330. eprint: https://doi.org/10.1021/acs.chemrev.6b00729. https://doi.org/10.1021/acs.chemrev.6b00729 (2017).

5. Von Diezmann, A., Shechtman, Y. & Moerner, W. Three-Dimensional Localization of Single Molecules for Super-Resolution Imaging and Single-Particle Tracking. Chem. Rev. 117. PMID: 28151646, 7244–7275. eprint: https://doi.org/10.1021/acs.chemrev.6b00629. https://doi.org/10.1021/acs.chemrev.6b00629 (2017).

6. Beghin, A. et al. Localization-based super-resolution imaging meets high-content screening. Nat. Methods 14, 1184–1190. issn: 1548-7105. https://doi.org/10.1038/nmeth.4486 (Dec. 2017).

7. Patterson, G., Davidson, M., Manley, S. & Lippincott-Schwartz, J. Superresolution Imaging using Single-Molecule Localization. Annu. Rev. Phys. Chem. 61. PMID: 20055680, 345–367. eprint: https://doi.org/10.1146/annurev.physchem.012809.103444. https://doi.org/10.1146/annurev.physchem.012809.103444 (2010).

8. Egner, A. et al. Fluorescence Nanoscopy in Whole Cells by Asynchronous Localization of Photoswitching Emitters. Biophys. J. 93, 3285–3290. issn: 0006-3495. https://www.sciencedirect.com/science/article/pii/S0006349507715825 (2007).

9. Thompson, R. E., Larson, D. R. & Webb, W. W. Precise nanometer localization analysis for individual fluorescent probes. Biophys. J. 82, 2775–2783. issn: 0006-3495. https://www.sciencedirect.com/science/article/pii/S000634950275618X (2002).

10. Dempsey, G. T., Vaughan, J. C., Chen, K. H., Bates, M. & Zhuang, X. Evaluation of fluorophores for optimal performance in localization-based super-resolution imaging. Nat. Methods 8, 1027–1036 (2011).

11. Hell, S. W. & Wichmann, J. Breaking the diffraction resolution limit by stimulated emission: stimulated-emission-depletion fluorescence microscopy. Opt. Lett. 19, 780–782. https://opg.optica.org/ol/abstract.cfm?URI=ol-19-11-780 (June 1994).

12. Huppa, J. B. & Schütz, G. J. in Encyclopedia of Cell Biology (Second Edition) (eds Bradshaw, R. A., Hart, G. W. & Stahl, P. D.) Second Edition, 536–552 (Academic Press, Oxford, 2023). isbn: 978-0-12-821624-8. https://www.sciencedirect.com/science/article/pii/B9780128216187002030.

13. Yang, L., Chen, Q., Wang, Z., Zhang, H. & Sun, H. Small-molecule fluorescent probes for plasma membrane staining: Design, mechanisms and biological applications. Coord. Chem. Rev. 474, 214862. issn: 0010-8545. https://www.sciencedirect.com/science/article/pii/S001085452200457X (2023).

14. Chen, K. H., Boettiger, A. N., Moffitt, J. R., Wang, S. & Zhuang, X. Spatially resolved, highly multiplexed RNA profiling in single cells. Science 348, aaa6090 (2015).

15. Singer, R. H. & Ward, D. C. Actin gene expression visualized in chicken muscle tissue culture by using in situ hybridization with a biotinated nucleotide analog. Proc. Natl. Acad. Sci. U.S.A. 79, 7331–7335 (1982).

16. Kilic, Z., Schweiger, M., Moyer, C., Shepherd, D. & Pressé, S. Gene expression model inference from snapshot RNA data using Bayesian non-parametrics. Nat. Comput. Sci., 1–10 (2023).

17. Fernández-Suárez, M. & Ting, A. Fluorescent probes for super-resolution imaging in living cells. Nat. Rev. Mol. Cell Biol. 9, 929–943. issn: 1471-0080. https://doi.org/10.1038/nrm2531 (Dec. 2008).

18. Gahlmann, A. & Moerner, W. Exploring bacterial cell biology with single-molecule tracking and super-resolution imaging. Nat. Rev. Microbiol. 12, 9–22 (2014).

19. Manley, S. et al. High-density mapping of single-molecule trajectories with photoactivated localization microscopy. Nat. Methods 5, 155–157 (2008).

20. Lambert, T. J. FPbase: a community-editable fluorescent protein database. Nat. Methods 16, 277–278. issn: 1548-7105. https://doi.org/10.1038/s41592-019-0352-8 (Apr. 2019).

21. Nixon, M. & Aguado, A. Feature extraction and image processing for computer vision(Academic press, 2019).

22. Chenouard, N. et al. Objective comparison of particle tracking methods. Nat. Methods 11, 281–289. issn: 1548-7105. https://doi.org/10.1038/nmeth.2808 (Mar. 2014).

23. Anderson, C. M., Georgiou, G. N., Morrison, I., Stevenson, G. & Cherry, R. J. Tracking of cell surface receptors by fluorescence digital imaging microscopy using a charge-coupled device camera. Low-density lipoprotein and influenza virus receptor mobility at 4 degrees C. J. Cell. Sci. 101, 415–425 (1992).

24. Seisenberger, G. et al. Real-time single-molecule imaging of the infection pathway of an adeno-associated virus. Science 294, 1929–1932 (2001).

25. Jaqaman, K. et al. Robust single-particle tracking in live-cell time-lapse sequences. Nat. Methods 5, 695–702. issn: 1548-7105. https://doi.org/10.1038/nmeth.1237 (Aug. 2008).

26. Lowe, D. G. Distinctive image features from scale-invariant keypoints. Int. J. Comput. Vis. 60, 91–110 (2004).

27. Vallotton, P., Ponti, A., Waterman-Storer, C., Salmon, E. & Danuser, G. Recovery, Visualization, and Analysis of Actin and Tubulin Polymer Flow in Live Cells: A Fluorescent Speckle Microscopy Study. Biophys. J. 85, 1289–1306. issn: 0006-3495. https://www.sciencedirect.com/science/article/pii/S0006349503745640 (2003).

28. Bonneau, S., Dahan, M. & Cohen, L. Single quantum dot tracking based on perceptual grouping using minimal paths in a spatiotemporal volume. IEEE Trans. Image Process. 14, 1384–1395 (2005).

29. Sbalzarini, I. & Koumoutsakos, P. Feature point tracking and trajectory analysis for video imaging in cell biology. J. Struct. Biol. 151, 182–195. issn: 1047-8477. https://www.sciencedirect.com/science/article/pii/S1047847705001267 (2005).

30. Carter, B. C., Shubeita, G. T. & Gross, S. P. Tracking single particles: A user-friendly quantitative evaluation. Phys. Biol. 2, 60–72. https://dx.doi.org/10.1088/1478-3967/2/1/008 (Mar. 2005).

31. Racine, V. et al. Multiple-target tracking of 3D fluorescent objects based on simulated annealing in 3rd IEEE International Symposium on Biomedical Imaging: Nano to Macro, 2006. (2006), 1020–1023.

32. Genovesio, A. et al. Multiple Particle Tracking in 3-D+t Microscopy: Method and Application to the Tracking of Endocytosed Quantum Dots. IEEE Trans. Image Process. 15, 1062–1070. issn: 1057-7149 (2006).

33. Fox, E., Sudderth, E. & Willsky, A. Hierarchical Dirichlet processes for tracking maneuvering targets in 2007 10th International Conference on Information Fusion (2007).

34. Smal, I., Draegestein, K., Galjart, N., Niessen, W. & Meijering, E. Particle filtering for multiple object tracking in dynamic fluorescence microscopy images: Application to microtubule growth analysis. 27, 789–804 (2008).

35. Sergé, A., Bertaux, N., Rigneault, H. & Marguet, D. Dynamic multiple-target tracing to probe spatiotemporal cartography of cell membranes. Nat. Methods 5, 687–694. issn: 1548-7105. https://doi.org/10.1038/nmeth.1233 (Aug. 2008).

36. Chenouard, N., Bloch, s. & Olivo-Marin, J.-C. Feature-aided particle tracking in 2008 15th IEEE International Conference on Image Processing (2008), 1796–1799.

37. Coraluppi, S. & Carthel, C. Multi-Stage Multiple-Hypothesis Tracking. J. Adv. Inf. Fusion 6, 57–67 (2011).

38. Coraluppi, S. & Carthel, C. Recursive track fusion for multi-sensor surveillance. Inf. Fusion 5, 23–33 (2004).

39. Olivo-Marin, J.-C. Extraction of spots in biological images using multiscale products. Pattern Recognit. 35, 1989–1996 (2002).

40. Winter, M. R., Fang, C., Banker, G., Roysam, B. & Cohen, A. R. Axonal transport analysis using multitemporal association tracking. Int. J. Comput. Biol. Drug Des 5, 35–48 (2012).

41. Winter, M. et al. Vertebrate neural stem cell segmentation, tracking and lineaging with validation and editing. Nat. Protoc. 6, 1942–1952 (2011).

42. Godinez, W. J., Lampe, M., Eils, R., Müller, B. & Rohr, K. Tracking multiple particles in fluorescence microscopy images via probabilistic data association in 2011 IEEE International Symposium on Biomedical Imaging: From Nano to Macro (2011), 1925–1928.

43. Rink, J., Ghigo, E., Kalaidzidis, Y. & Zerial, M. Rab conversion as a mechanism of progression from early to late endosomes. Cell 122, 735–749 (2005).

44. Liang, L., Shen, H., De Camilli, P. & Duncan, J. S. Tracking clathrin coated pits with a multiple hypothesis based method in Medical Image Computing and Computer-Assisted Intervention–MICCAI 2010: 13th International Conference, Beijing, China, September 20-24, 2010, Proceedings, Part II 13 (2010), 315–322.

45. Magnusson, K. E. & Jaldén, J. A batch algorithm using iterative application of the Viterbi algorithm to track cells and construct cell lineages in 2012 9th IEEE International Symposium on Biomedical Imaging (ISBI) (2012), 382–385.

46. Crocker, J. C. & Grier, D. G. Methods of digital video microscopy for colloidal studies. J. Colloid Interface Sci. 179, 298–310 (1996).

47. Casuso, I. et al. Characterization of the motion of membrane proteins using high-speed atomic force microscopy. Nat. Nanotechnol. 7, 525–529 (2012).

48. Husain, M., Boudier, T., Paul-Gilloteaux, P., Casuso, I. & Scheuring, S. Software for drift compensation, particle tracking and particle analysis of high-speed atomic force microscopy image series. J. Mol. Recognit. 25, 292–298 (2012).

49. Rousseeuw, P. J. & Leroy, A. M. Robust regression and outlier detection (John wiley & sons, 2005).

50. Shafique, K. & Shah, M. A noniterative greedy algorithm for multiframe point correspondence. IEEE Trans. Pattern Anal. Mach. Intell. 27, 51–65 (2005).

51. Ku, T.-C., Kao, L.-S., Lin, C.-C. & Tsai, Y.-S. Morphological filter improve the efficiency of automated tracking of secretory vesicles with various dynamic properties. Microsc. Res. Tech. 72, 639–649 (2009).

52. Ku, T.-C. et al. An automated tracking system to measure the dynamic properties of vesicles in living cells. Microsc. Res. Tech. 70, 119–134 (2007).

53. Celler, K., van Wezel, G. P. & Willemse, J. Single particle tracking of dynamically localizing TatA complexes in Streptomyces coelicolor. Biochem. Biophys. Res. Commun. 438, 38–42 (2013).

54. Godinez, W. et al. Deterministic and Probabilistic Approaches for Tracking Virus Particles in Time-Lapse Fluorescence Microscopy Image Sequences. Med. Image Anal. 13, 325–342. issn: 1361-8415. https://www.sciencedirect.com/science/article/pii/S1361841508001412 (2009).

55. Xue, Q. & Leake, M. C. A novel multiple particle tracking algorithm for noisy in vivo data by minimal path optimization within the spatio-temporal volume in 2009 IEEE International Symposium on Biomedical Imaging: From Nano to Macro (2009), 1158–1161.

56. Chenouard, N., Dufour, A. & Olivo-Marin, J.-C. Tracking algorithms chase down pathogens. Biotechnol. J. 4, 838–845. eprint: https://onlinelibrary.wiley.com/doi/pdf/10.1002/biot.200900030. https://onlinelibrary.wiley.com/doi/abs/10.1002/biot.200900030 (2009).

57. Chertkov, M., Kroc, L., Krzakala, F., Vergassola, M. & Zdeborová, L. Inference in particle tracking experiments by passing messages between images. Proc. Natl. Acad. Sci. U.S.A. 107, 7663–7668. eprint: https://www.pnas.org/doi/pdf/10.1073/pnas.0910994107. https://www.pnas.org/doi/abs/10.1073/pnas.0910994107 (2010).

58. Agrawal, A., Gupta, M., Veeraraghavan, A. & Narasimhan, S. G. Optimal coded sampling for temporal super-resolution in 2010 IEEE Computer Society Conference on Computer Vision and Pattern Recognition (2010), 599–606.

59. Park, H. Y., Buxbaum, A. R. & Singer, R. H. in Single Molecule Tools: Fluorescence Based Approaches, Part A (ed Walter, N. G.) 387–406 (Academic Press, 2010). https://www.sciencedirect.com/science/article/pii/S0076687910720036.

60. Dufour, A., Thibeaux, R., Labruyere, E., Guillen, N. & Olivo-Marin, J.-C. 3-D Active Meshes: Fast Discrete Deformable Models for Cell Tracking in 3-D Time-Lapse Microscopy. IEEE Trans. Image Process. 20, 1925–1937 (2011).

61. Mudenagudi, U., Banerjee, S. & Kalra, P. K. Space-Time Super-Resolution Using Graph-Cut Optimization. IEEE Transactions on Pattern Analysis and Machine Intelligence 33, 995–1008 (2011).

62. Meijering, E., Dzyubachyk, O. & Smal, I. in Imaging and Spectroscopic Analysis of Living Cells (ed conn, P. M.) 183–200 (Academic Press, 2012). https://www.sciencedirect.com/science/article/pii/B9780123918574000094.

63. Michalet, X. & Berglund, A. Optimal Diffusion Coefficient Estimation in Single-Particle Tracking. Phys. Rev. E 85, 061916. issn: 1539-3755. https://link.aps.org/doi/10.1103/PhysRevE.85.061916 (6 June 2012).

64. Chenouard, N., Bloch, I. & Olivo-Marin, J.-C. Multiple Hypothesis Tracking for Cluttered Biological Image Sequences. IEEE Trans. Pattern Anal. Mach. Intell. 35, 2736–3750 (2013).

65. Persson, F., Lindén, M., Unoson, C. & Elf, J. Extracting intracellular diffusive states and transition rates from single-molecule tracking data. Nat. Methods 10, 265–269. issn: 1548-7105. https://doi.org/10.1038/nmeth.2367 (Mar. 2013).

66. Mont, A. D., Calderon, C. P. & Poore, A. B. A new computational method for ambiguity assessment of solutions to assignment problems arising in target tracking in Signal and Data Processing of Small Targets 2014 (ed Drummond, O. E.) 9092 (SPIE, 2014), 159–176. https://doi.org/10.1117/12.2053400.

67. Rowland, D. J. & Biteen, J. S. Top-Hat and Asymmetric Gaussian-Based Fitting Functions for Quantifying Directional Single-Molecule Motion. ChemPhysChem 15, 712–720. eprint: https://chemistry-europe.onlinelibrary.wiley.com/doi/pdf/10.1002/cphc.201300774. https://chemistry-europe.onlinelibrary.wiley.com/doi/abs/10.1002/cphc.201300774 (2014).

68. Fox, E. B., Hughes, M. C., Sudderth, E. B. & Jordan, M. I. Joint Modeling of Multiple Time Series via the Beta Process with Application to Motion Capture Segmentation. Ann. Appl. Stat. 8, 1281–1313. https://doi.org/10.1214/14-AOAS742 (2014).

69. Barden, A. O. et al. Tracking individual membrane proteins and their biochemistry: The power of direct observation. Neuropharmacology 98. Fluorescent Tools in Neuropharmacology, 22–30. issn: 0028-3908. https://www.sciencedirect.com/science/article/pii/S0028390815001847 (2015).

70. Monnier, N. et al. Inferring transient particle transport dynamics in live cells. Nat. Methods 12, 838–840. issn: 1548-7105. https://doi.org/10.1038/nmeth.3483 (Sept. 2015).

71. Maroulas, V. & Nebenführ, A. Tracking rapid intracellular movements: A Bayesian random set approach. Ann. Appl. Stat. 9, 926–949. https://doi.org/10.1214/15-AOAS819 (2015).

72. Tinevez, J.-Y. et al. TrackMate: An open and extensible platform for single-particle tracking. Methods 115. Image Processing for Biologists, 80–90. issn: 046-2023. https://www.sciencedirect.com/science/article/pii/S1046202316303346 (2017).

73. Sgouralis, I., Nebenführ, A. & Maroulas, V. A Bayesian Topological Framework for the Identification and Reconstruction of Subcellular Motion. SIAM J. Imaging Sci. 10, 871–899. eprint: https://doi.org/10.1137/16M1095755. https://doi.org/10.1137/16M1095755 (2017).

74. Saxton, M. J. & Jacobson, K. SINGLE-PARTICLE TRACKING:Applications to Membrane Dynamics. Annu. Rev. Biophys. 26. PMID: 9241424, 373–399. eprint: https://doi.org/10.1146/annurev.biophys.26.1.373. https://doi.org/10.1146/annurev.biophys.26.1.373 (1997).

75. Cheng, H.-J., Hsu, C.-H., Hung, C.-L. & Lin, C.-Y. A review for cell and particle tracking on microscopy images using algorithms and deep learning technologies. Biomed. J. 45, 465–471. issn: 2319-4170. https://www.sciencedirect.com/science/article/pii/S2319417021001359 (2022).

76. De Chaumont, F. et al. Icy: an open bioimage informatics platform for extended reproducible research. Nat. Methods 9, 690–696. issn: 1548-7105. https://doi.org/10.1038/nmeth.2075 (July 2012).

77. Tameling, C. et al. Colocalization for super-resolution microscopy via optimal transport. Nat. Comput. Sci 1, 199–211. issn: 2662-8457. https://doi.org/10.1038/s43588-021-00050-x (Mar. 2021).

78. Ladouceur, A.-M. et al. Clusters of bacterial RNA polymerase are biomolecular condensates that assemble through liquid-liquid phase separation. Proc. Natl. Acad. Sci. U.S.A. 117, 18540–18549. eprint: https://www.pnas.org/doi/pdf/10.1073/pnas.2005019117. https://www.pnas.org/doi/abs/10.1073/pnas.2005019117 (2020).

79. Lim, D., Ford, T. N., Chu, K. K. & Mertz, J. Optically sectioned in vivo imaging with speckle illumination HiLo microscopy. J. Biomed. Opt. 16, 016014–016014 (2011).

80. Smith, C. S., Joseph, N., Rieger, B. & Lidke, K. A. Fast, single-molecule localization that achieves theoretically minimum uncertainty. Nat. Methods 7, 373–375 (2010).

81. Abraham, A. V., Ram, S., Chao, J., Ward, E. & Ober, R. J. Quantitative study of single molecule location estimation techniques. Opt. Express 17, 23352–23373 (2009).

82. Huang, Z.-L. et al. Localization-based super-resolution microscopy with an sCMOS camera. Opt. Express 19, 19156–19168 (2011).

83. Zhang, B., Zerubia, J. & Olivo-Marin, J.-C. Gaussian approximations of fluorescence microscope point-spread function models. Appl. Opt. 46, 1819–1829 (2007).

84. Geman, S. & Geman, D. Stochastic Relaxation, Gibbs Distributions, and the Bayesian Restoration of Images. IEEE Transactions on Pattern Analysis and Machine Intelligence PAMI-6, 721–741 (1984).

85. Metropolis, N., Rosenbluth, A. W., Rosenbluth, M. N., Teller, A. H. & Teller, E. Equation of State Calculations by Fast Computing Machines. J. Chem. Phys. 21, 1087–1092. eprint: https://doi.org/10.1063/1.1699114. https://doi.org/10.1063/1.1699114 (1953).

86. Hastings, W. K. Monte Carlo Sampling Methods Using Markov Chains and Their Applications. Biometrika 57, 97–109. issn: 00063444. http://www.jstor.org/stable/2334940 (1970).

87. Danisch, S. & Krumbiegel, J. Makie.jl: Flexible high-performance data visualization for Julia. J. Open Source Softw. 6, 3349. https://doi.org/10.21105/joss.03349 (2021).

88. Bezanson, J., Edelman, A., Karpinski, S. & Shah, V. B. Julia: A fresh approach to numerical computing. SIAM review 59, 65–98. https://doi.org/10.1137/141000671 (2017).

89. Schindelin, J. et al. Fiji: an open-source platform for biological-image analysis. Nat. Methods 9, 676–682. issn: 1548-7105. https://doi.org/10.1038/nmeth.2019 (July 2012).

