## Supplemental Materials for "BNP-Track: A framework for multi-particle superresolved tracking"

Lance W.Q. Xu (徐伟青)[*1,2], Ioannis Sgouralis[*3], Zeliha Kilic[4], and Steve Pressé [†1,2,5]

[*]Contributed equally.

### 1 Additional tables

| Description | Parameter | Unit | Reference value |
|---|---|---|---|
| Background flux | $F$ | $\mu m^{-2}s^{-1}$ | $10^5$ |
| Diffusion coefficient | $D$ | $\mu m^2 s^{-1}$ | 0.05 |
| Emission rate | $h$ | $s^{-1}$ | $10^4$ |
| Emitter number | $B$ | | 1 |
| Gain | $G$ | $ADU^{-1}$ | 0.3 |

Table S.1: List of symbols and their reference values.

| Supplementary video | $B$ | $D$ | $h$ | $F$ | Appear in |
|---|---|---|---|---|---|
| 1 | 1 | 0.05 | $10^4$ | $10^5$ | Figs. S.1a, 2, 4a and 4b, Figs. S.3a to S.3d |
| 2 | 3 | 0.05 | $10^4$ | $10^5$ | Figs. S.1b, 3, 4c, 4d and 6c to 6e, Figs. S.3e to S.3h, Figs. S.6d to S.6f, S.8a, S.8b, S.9 and S.10 |
| 3 | 3 | 0.05 | $10^4$ | $10^5$ | Fig. S.2 |
| 4 | 7 | 0.05 | $10^4$ | $10^5$ | Figs. S.3i to S.3l |
| 5 | 1 | 0.05 | $10^4$ | $10^5$ | Fig. 5 |
| 6 | 0.01 | 3 | $10^4$ | $10^5$ | Figs. 6a to 6c |
| 7 | 0.25 | 3 | $10^4$ | $10^5$ | Figs. 6g to 6i |
| 8 | 1 | 1 | $10^4$ | $10^5$ | Fig. S.5 |
| 9 | 3 | 0.05 | $2 \times 10^3$ | $10^5$ | Figs. S.6a to S.6c |
| 10 | 3 | 0.05 | $5 \times 10^4$ | $10^5$ | Figs. S.6g to S.6i |
| 11 | 3 | 0.05 | $10^4$ | $2 \times 10^4$ | Figs. S.7a to S.7c |
| 12 | 3 | 0.05 | $10^4$ | $5 \times 10^5$ | Figs. S.7g to S.7i |
| 13 | 3 | 0.05 | $10^4$ | $10^5$ | Figs. S.8c and S.8d |

Supplementary Video 3 contains emitters that are closer to each other than Supplementary video 2.

Supplementary Video 13 contains an out-of-focus emitter that is roughly 500 nm away from the in focus plane.

Table S.2: The index of all Supplementary videos. Each symbol's definition is in Table S.1.

| Description | Parameter | Unit | Value |
|---|---|---|---|
| Emission wavelength | $\lambda$ | nm | 665 |
| Exposure period | $\tau$ | s | 0.03 |
| Frame period | | s | 0.033 |
| Image height | | pixel | 23 |
| Image with | | pixel | 15 |
| Numerical aperture | NA | | 1.45 |
| Pixel size | | nm | 133 |
| Refractive index | $n_{\mathrm{RI}}$ | | 1.515 |
| Noise excess factor | $f$ | | 2 |
| Quantum efficiency | $\beta$ | | |

Table S.3: List of camera parameter values and their symbols.

| | BNP-Track | TrackMate A | u-track |
|---|---|---|---|
| Global measures | | | |
| Pairing distance | 13.0 | 13.3 | 12.1 |
| Normalized pairing score (alpha) | 0.948 | 0.947 | 0.952 |
| Full normalized score (beta) | 0.948 | 0.947 | 0.952 |
| Tracks | | | |
| Number of reference tracks | 1 | 1 | 1 |
| Number of candidate tracks | 1 | 1 | 1 |
| Similarity between tracks (Jaccard) | 1.0 | 1.0 | 1.0 |
| Number of paired tracks | 1 | 1 | 1 |
| Number of missed tracks | 0 | 0 | 0 |
| Number of spurious tracks | 0 | 0 | 0 |
| Detections | | | |
| Number of reference detections | 50 | 50 | 50 |
| Number of candidate detections | 50 | 50 | 50 |
| Similarity between detections (Jaccard) | 1.0 | 1.0 | 1.0 |
| Number of paired detections | 50 | 50 | 50 |
| Number of missed detections | 0 | 0 | 0 |
| Number of spurious detections | 0 | 0 | 0 |
| Detection accuracy | | | |
| Root mean-square error | 0.300 | 0.314 | 0.297 |
| Minimum distance | 0.0310 | 0.0168 | 0.0241 |
| Maximum distance | 0.660 | 0.809 | 0.888 |
| Distance standard deviation | 0.150 | 0.166 | 0.173 |

Table S.4: Full tracking performance measure comparison among BNP-Track, TrackMate, and u-track for Supplementary Video 1.

| | BNP-Track | TrackMate A | TrackMate B |
|---|---|---|---|
| Global measures | | | |
| Pairing distance | 75.4 | 337 | 386 |
| Normalized pairing score (alpha) | 0.899 | 0.550 | 0.484 |
| Full normalized score (beta) | 0.899 | 0.550 | 0.484 |
| Tracks | | | |
| Number of reference tracks | 3 | 3 | 3 |
| Number of candidate tracks | 3 | 3 | 3 |
| Similarity between tracks (Jaccard) | 1 | 1 | 1 |
| Number of paired tracks | 3 | 3 | 3 |
| Number of missed tracks | 0 | 0 | 0 |
| Number of spurious tracks | 0 | 0 | 0 |
| Detections | | | |
| Number of reference detections | 150 | 150 | 150 |
| Number of candidate detections | 150 | 106 | 93 |
| Similarity between detections (Jaccard) | 1.0 | 0.593 | 0.62 |
| Number of paired detections | 150 | 89 | 93 |
| Number of missed detections | 0 | 61 | 57 |
| Number of spurious detections | 0 | 0 | 1 |
| Detection accuracy | | | |
| Root mean-square error | 0.807 | 0.473 | 1.77 |
| Minimum distance | 0.0441 | 0.0242 | 0.0227 |
| Maximum distance | 3.62 | 1.07 | 4.65 |
| Distance standard deviation | 0.631 | 0.303 | 1.40 |

Table S.5: Full tracking performance measure comparison between BNP-Track and TrackMate for the three-emitter video.

#### 2 Additional figures

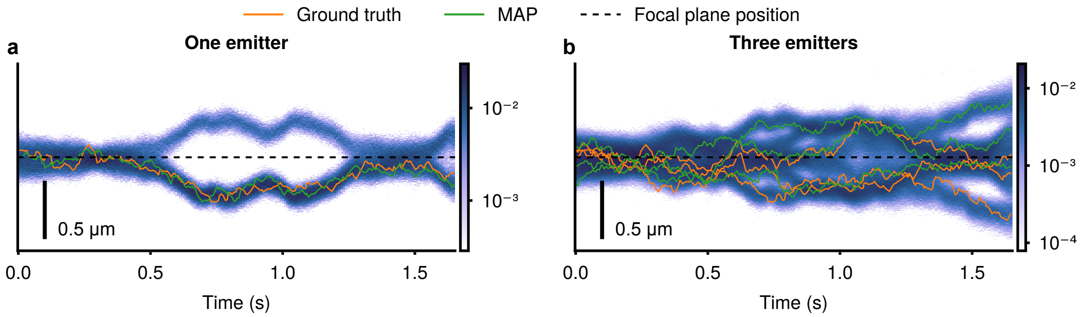

Figure S.1: BNP-Track's performance in the determination of axial position, *i.e.*, $z$ trajectories, in the one-emitter case (Supplementary Video 1), **a**, and the three-emitter case (Supplementary Video 2), **b**. As the PSF used in this study is symmetric about the in focus plane, we cannot distinguish between a location and its reflection about the plane. This explains why our estimates are symmetric about the in focus plane (marked as a black dashed line).

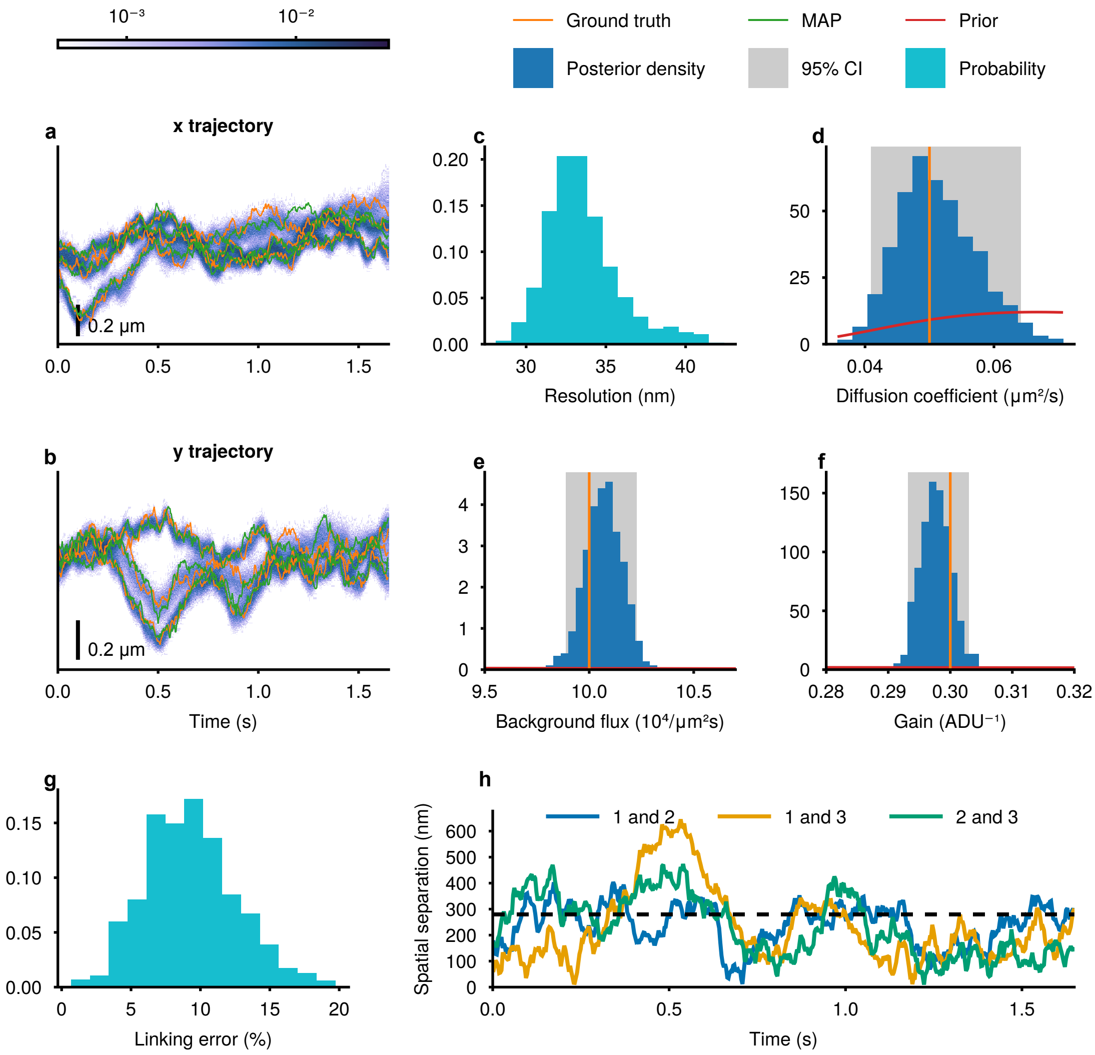

Figure S.2: A similar plot as in Fig. 3 but with a video (Supplementary Video 3) where emitters are even closer to each other in space. BNP-Track can still estimate the correct number of emitters as well as other parameters of interest, but with higher mean localization and linking error. This dataset is provided as Supplementary Video 3.

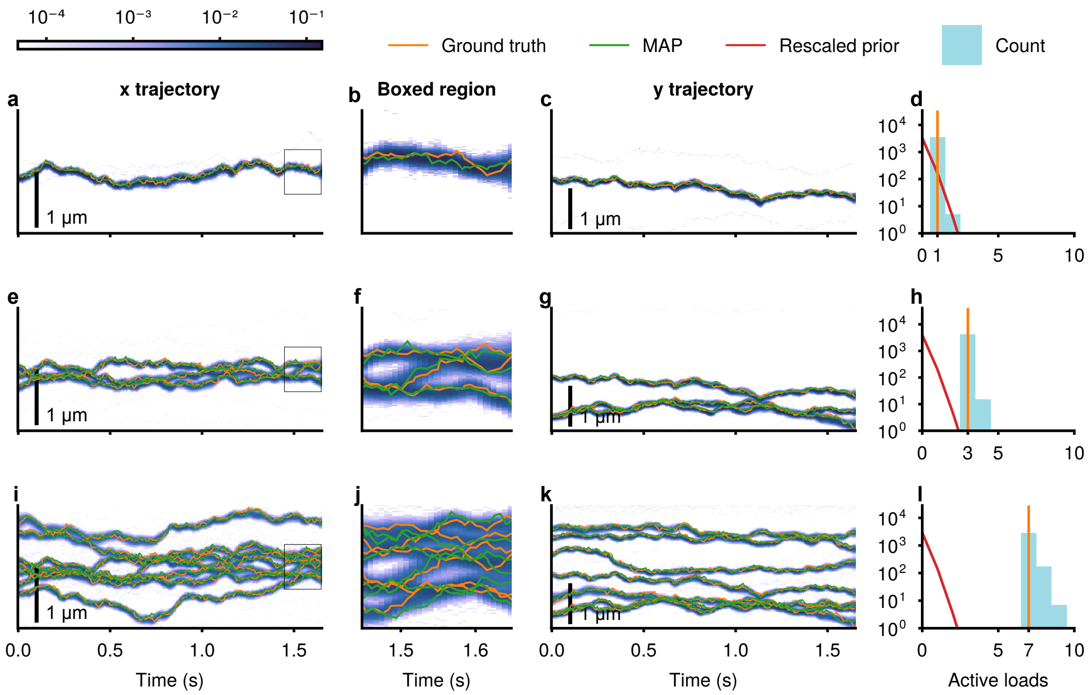

Figure S.3: BNP-Track's performance with different emitter numbers. **a-d** are from Supplementary Video 1 with one emitter, **e-h** are from Supplementary Video 2 with three emitters, and **i-l** are from Supplementary Video 4 with seven emitters.

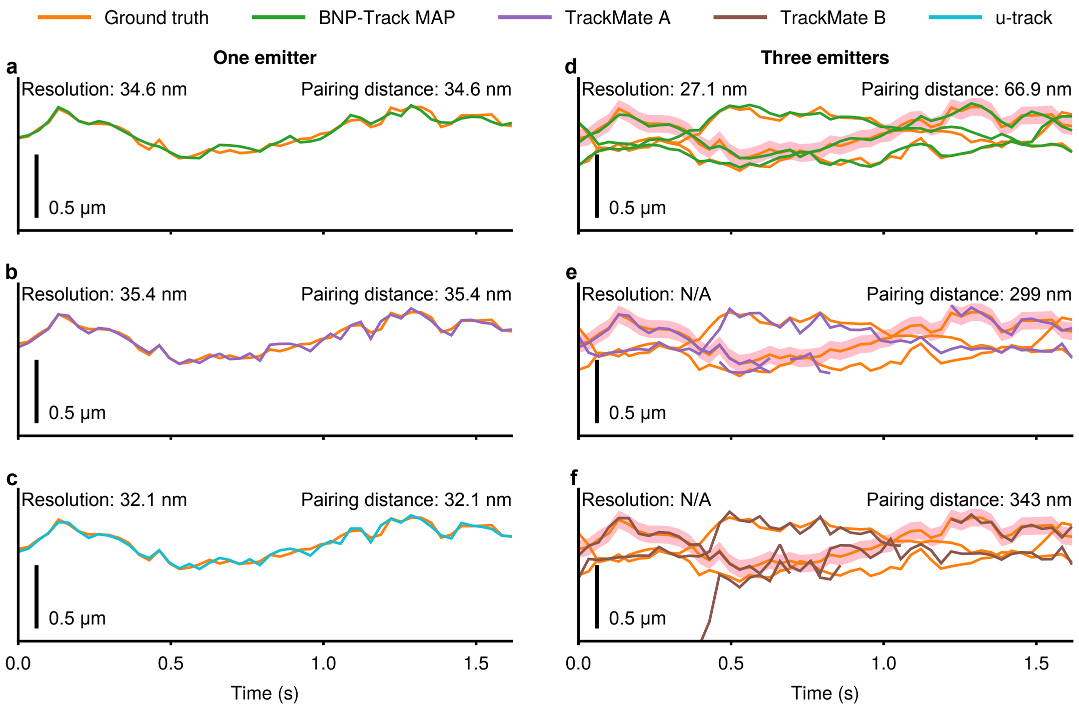

Figure S.4: Same layout as Fig. 4 but for the $x$ coordinate.

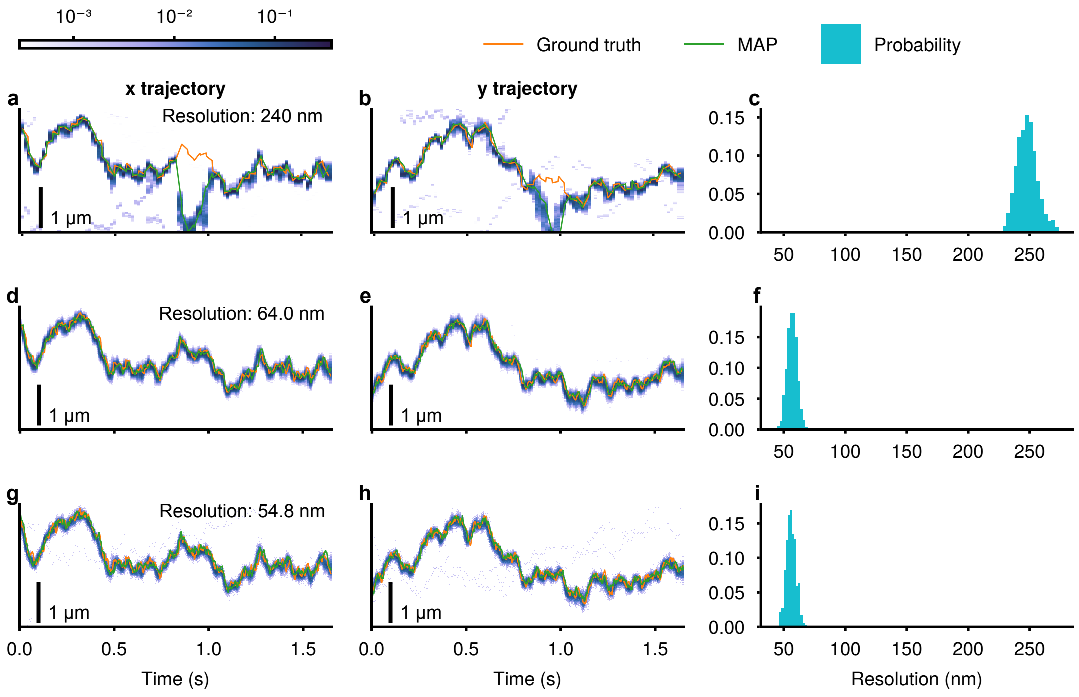

Figure S.5: BNP-Track's performance testing the number of positions sampled within the exposure period of one frame. This synthetic video (Supplementary Video 8) has one emitter diffusing at $1\,\mu\mathrm{m}^2\mathrm{s}^{-1}$. In **a**-**c** BNP-Track only samples two positions within each exposure, while in **d**-**f** there are five, and in **g**-**i** there are ten. For this faster diffusing case, inferring two position per frame fails to correctly track the emitter in some frames, yielding a poor localization resolution, see (**c**). Once the number of positions sampled within one exposure is sufficient (greater than or equal to five in this case), increasing this number will no longer improve the resolution, see (**f**) and (**i**).

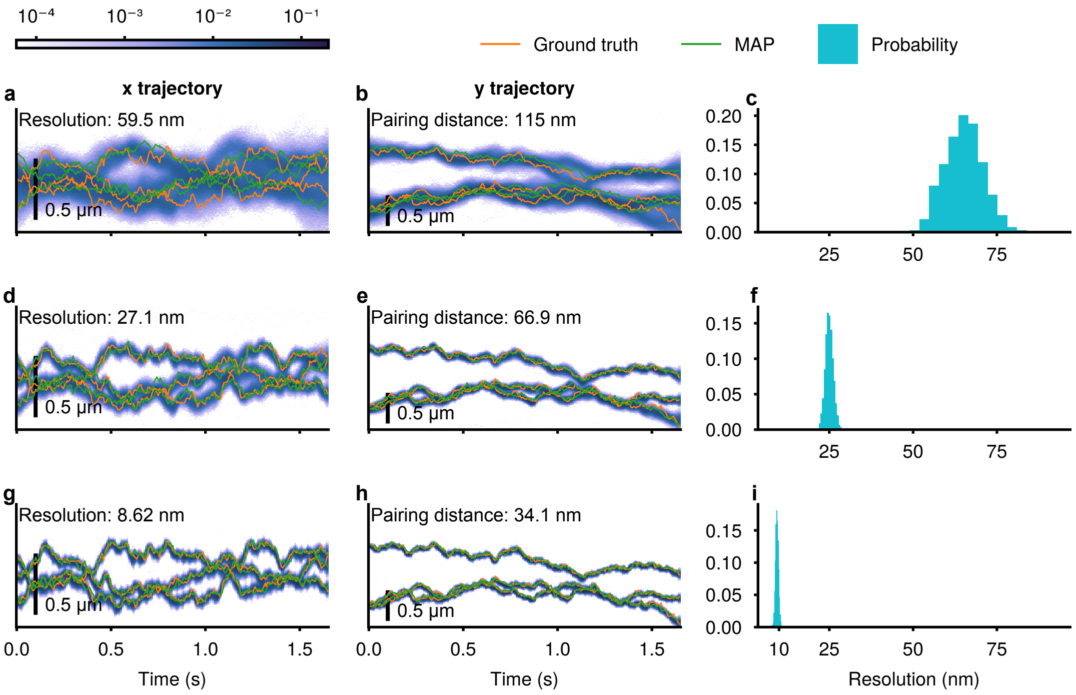

Figure S.6: BNP-Track's performance regarding different emission rates. **a**-**c** are from Supplementary Video 9 with emission rate $2 \times 10^3\,\mathrm{s}^{-1}$, **d**-**f** are from Supplementary Video 2 with emission rate being $10^4\,\mathrm{s}^{-1}$, and **g**-**i** are from Supplementary Video 10 with emission rate $5 \times 10^4\,\mathrm{s}^{-1}$.

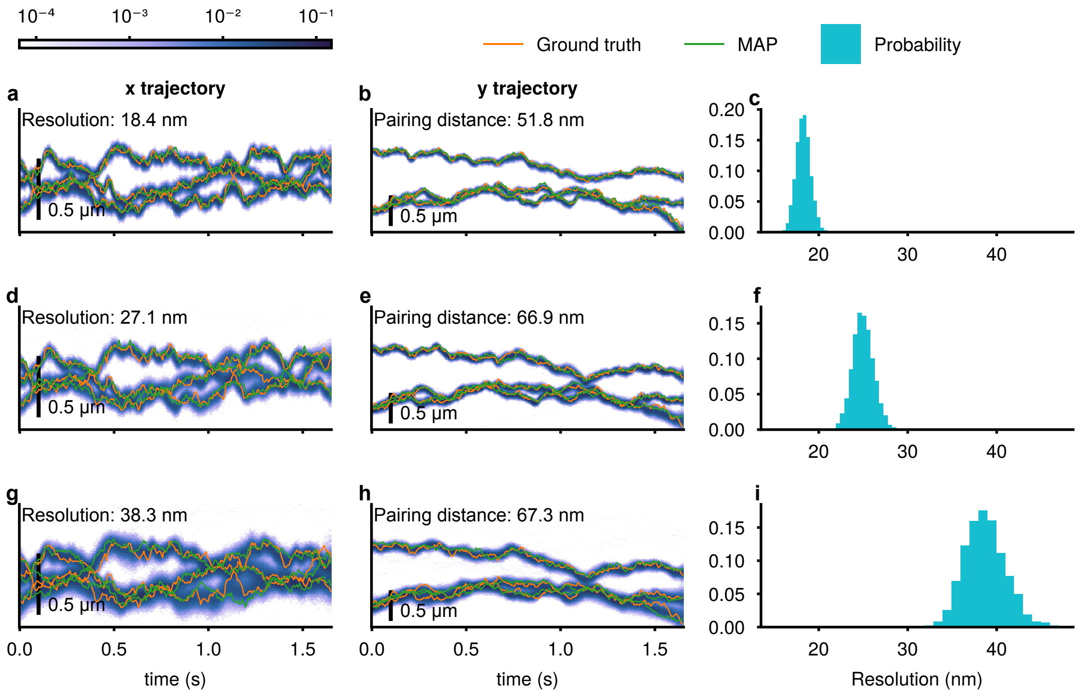

Figure S.7: BNP-Track's performance testing different background flux. **a-c** are from Supplementary Video 11 with background photon flux $2 \times 10^4\,\mu\text{m}^{-2}\text{s}^{-1}$, **d-f** are from Supplementary Video 2 with background photon flux $10^5\,\mu\text{m}^{-2}\text{s}^{-1}$, and **g-i** are from Supplementary Video 12 with background photon flux $5 \times 10^5\,\mu\text{m}^{-2}\text{s}^{-1}$.

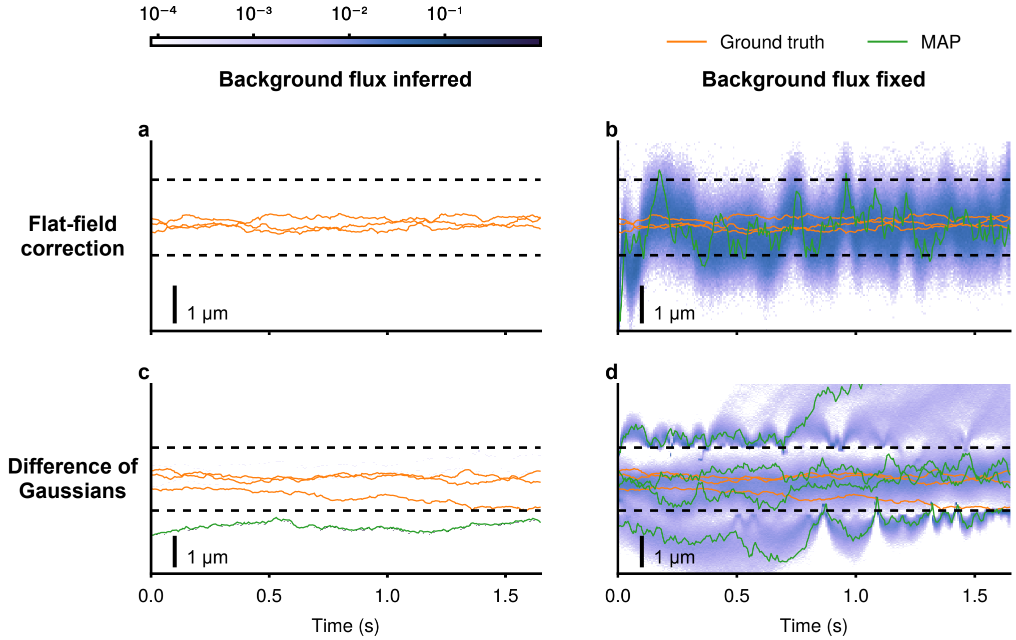

Figure S.8: BNP-Track's performance when used with naive background subtraction as is common in [1–5]. **a** and **b**, An video of pure background was simulated, and its temporal average was subtracted from each frame of Supplementary Video 2. BNP-Track can no longer infer any emitters while learning the background flux as each frame's mathematical model is now changed after the background subtraction. **c** and **d**, The same background subtraction as (**a**) and (**b**) are performed, but BNP-Track was run with background fixed at zero. **e** and **f**, A Gaussian filter (with standard deviation at four pixels) was applied to each frame of Supplementary Video 13 and the resulting image was taken as background.

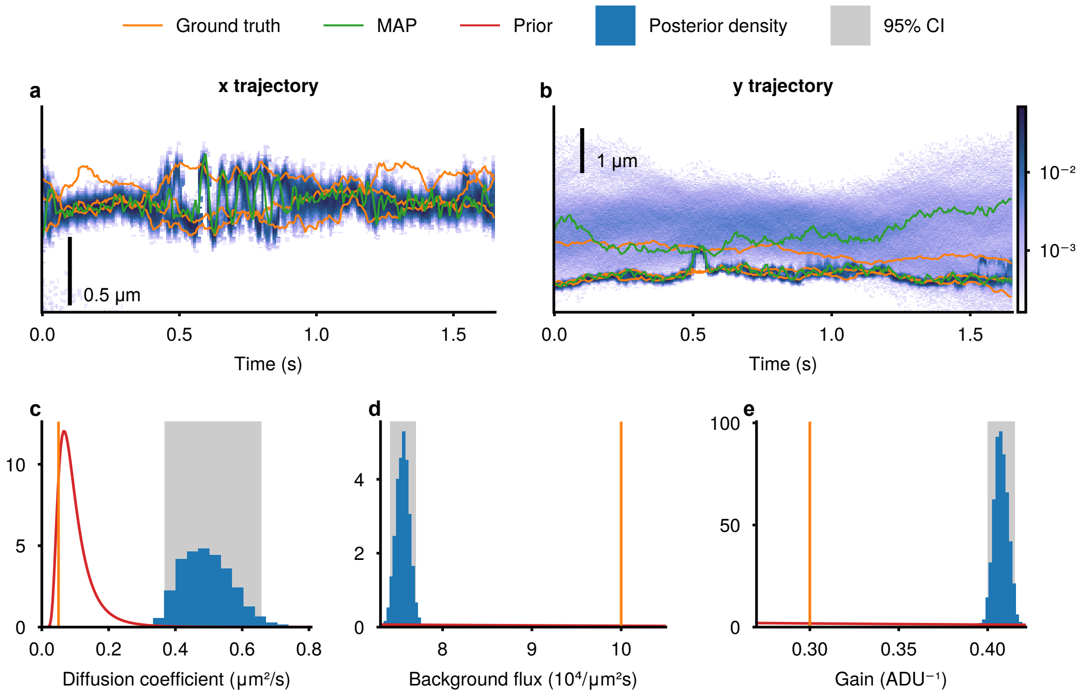

Figure S.9: BNP-Track's performance when the emitter number is fixed at two while there are three emitters (Supplementary Video 2). **a** and **b**, BNP-Track is forced to infer tracks jumping between actual trajectories. This results in incorrect estimates for all parameters, **c**-**e**, and highlights the importance of estimating all simultaneously and self-consistently.

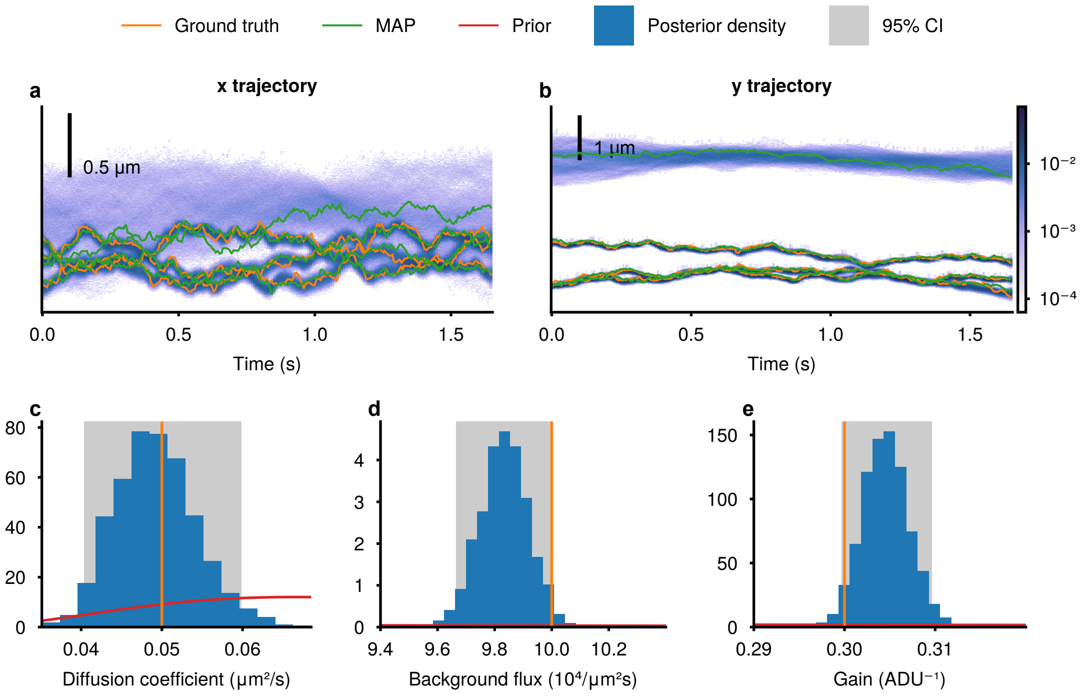

Figure S.10: BNP-Track's performance when the emitter number is fixed at four while there are three emitters (Supplementary Video 2). BNP-Track is able to infer correct tracks and parameters. The extra emitter is placed to be far away from the FOV of frames such that its contribution is negligible.

### 3 Forward model

#### C.1 Observation model

A key part of our work is modeling detectors (cameras) and this is critical, say, in correcting for motion blur biases (Figs. 1e to 1h), and re-assigning photons across pixels to the same emitter's PSF (*e.g.*, Eq. (6) where we integrate the PSF over the pixel area). For concreteness only, we focus on the electron-multiplying charge-coupled device (EMCCD) image detectors though different models, say for sCMOS, would amount to different expressions for Eq. (8) in main.

We denote the total number of frames by $N$ and the total number of pixels by $P$. We use the indices $n$ and $p$ to represent the $n$th frame and the $p$th pixel, respectively. At the $n$th frame and the $p$th pixel, the raw observation is denoted as $\underline{w}_n^p$, which is measured in terms of the analog-to-digital unit (ADU). We assume that each pixel has its own time-independent offset, $\mu^p$, which can be calibrated to obtain the corrected observation: $w_n^p = \underline{w}_n^p - \mu^p$.

We can use the following observation model, given the incident photon counts of the $p$th pixel within the $n$th frame ($u_n^p$), the gain of the sensor ($G$), the noise excess factor ($f$), and the quantum efficiency ($\beta$),

$$w_n^p | u_n^p, G \sim \mathbf{Gamma}\left(\frac{\beta u_n^p}{f}, fG\right). \tag{S.1}$$

This model states that "$w_n^p$ given $u_n^p$ and $G$ is sampled from a Gamma distribution with shape $\beta u_n^p f$ and scale $fG$". It is important to note that both $f$ and $\beta$ are treated as parameters that we can pre-calibrate; for EMCCDs, $f$ is equal to 2.

In order to model the incident photon counts $u_n^p$, we need to integrate the incident photon flux over the entire exposure time and pixel area. The exposure time interval in the $n$th frame is denoted as $[t_{n-1}, t_n]$, and the $p$th pixel is defined as the rectangular region $[x_{\min}^p, x_{\max}^p] \times [y_{\min}^p, y_{\max}^p]$. Therefore, the incident photon count in the $n$th frame and $p$th pixel can be represented by

$$u_n^p = \int_{x_{\min}^p}^{x_{\max}^p} \mathrm{d}x^p \int_{y_{\min}^p}^{y_{\max}^p} \mathrm{d}y^p \int_{t_{n-1}}^{t_n} \mathrm{d}t\, U(x^p, y^p, t), \tag{S.2}$$

where $U(x^p, y^p, t)$ is the incident photon flux with units of number per unit time per unit area. Additionally, the incident photon flux can be separated into two components: the background and actual emitter (fluorescently labeled molecules) contribution. Thus, we can represent the incident photon flux as shown,

$$U\left(x^p, y^p, t\right) = U_{\text{back}}\left(x^p, y^p, t\right) + U_{\text{fluor}}\left(x^p, y^p, t\right), \tag{S.3}$$

where $U_{\text{back}}\left(x^p, y^p, t\right)$ is the background photon flux and $U_{\text{fluor}}\left(x^p, y^p, t\right)$ is the fluorescence photon flux. We assume that the background photon flux is constant over both space and time, so the function $U_{\text{back}}(x^p, y^p, t)$ can be simplified to just $F$. As for the fluorescence photon flux $U_{\text{fluor}}$, we model it as the sum of the individual emitter contributions

$$U_{\text{fluor}}(x^p, y^p, t) = \sum_{m=1}^{M} b^m U_{\text{fluor}}^m(x^p, y^p; \mathbf{x}^m(t)), \tag{S.4}$$

where $M$ stands for the total number of active loads, $b^m \in \{0, 1\}$ is a label (called load) for whether the $m$th emitter is active, and $\mathbf{x}^m(t)$ is the spatial location of the $m$th emitter at time $t$. The individual emitter contribution is given by

$$U_{\text{fluor}}^m(x^p, y^p; \mathbf{x}^m(t)) = h\,\text{PSF}(x^p, y^p; \mathbf{x}^m(t)). \tag{S.5}$$

In this equation, $h$ is the emission rate or brightness of the emitter, assumed constant across all emitters. The point spread function (PSF), denoted as PSF $(x^p, y^p; \mathbf{x})$, satisfies the condition $\int_{-\infty}^{\infty}\int_{-\infty}^{\infty} \mathrm{d}x^p \mathrm{d}y^p \text{PSF}(x^p, y^p; \mathbf{x}^m(t)) = 1$. When these elements are plugged into the incident photon count formulation (Eq. (S.2)), we obtain

$$u_n^p = FA^p\tau + h\sum_{m=1}^{m} b^m \int_{t_{n-1}}^{t_n} \mathrm{d}t \int_{x_{\text{min}}^p}^{x_{\text{max}}^p} \mathrm{d}x^p \int_{y_{\text{min}}^p}^{y_{\text{max}}^p} \mathrm{d}y^p \text{PSF}(x^p, y^p; \mathbf{x}^m(t)). \tag{S.6}$$

Here, $A^p = (x_{\text{max}}^p - x_{\text{min}}^p) \times (y_{\text{max}}^p - y_{\text{min}}^p)$ represents the area of the $p$th pixel, $t_n$ is the time when the $n$th frame is produced, and $\tau$ is the exposure time of each frame. We note that $\tau \leq t_n - t_{n-1}$ due to camera dead time, and we assume no gaps between pixels, so $x_{\text{max}}^p = x_{\text{min}}^{p+1}$, $y_{\text{max}}^p = y_{\text{min}}^{p+1}$.

To evaluate the triple integral of Eq. (S.6), we must specify the integrand, namely the PSF. In practice, we can use any pre-calibrated form including forms induced by optical aberration. For concreteness here, we use the circular Gaussian Lorentzian PSF given by

$$\text{PSF}(x^p, y^p; \mathbf{x}^m(t)) = \frac{1}{2\pi\sigma_{z^m}^2} \exp\left(-\frac{(x^p - x^m)^2 + (y^p - y^m)^2}{2\sigma_{z^m}^2}\right), \tag{S.7}$$

$$\sigma_{z^m}^2 = \sigma_{\text{ref}}^2 \left[1 + \left(\frac{z^m}{Z_{\text{ref}}}\right)^2\right], \quad Z_{\text{ref}} = \frac{4\pi n_{\text{fluid}}}{\lambda}\sigma_{\text{ref}}^2. \tag{S.8}$$

The PSF's width, denoted by $\sigma_{z^m}$, is given in Eq. (S.8), where $n_{\text{fluid}}$ represents the refractive index of the immersion media in which the objective lens and the specimen are immersed, and $\lambda$ is the vacuum wave length of the emitted photons. The lateral Gaussian parameter $\sigma_{\text{ref}}$ is calculated differently based on the fluorescence microscope model [6]. In this work, we consider the nonparaxial widefield fluorescence microscope [7], and $\sigma_{\text{ref}}$ is given by

$$\sigma_{\text{ref}} = \frac{\lambda}{2\pi n_{\text{fluid}}} \sqrt{\frac{7\,(1 - \cos^{3/2}\alpha)}{7\,(4 - 7\cos^{3/2}\alpha + 3\cos^{7/2}\alpha)}} \tag{S.9}$$

where $\alpha$ is the maximal convergence semi-angle of the objective.

Now that the PSF has been specified, we can evaluate analytically the integral in Eq. (6). The area integral is then,

$$\int_{x^p_{\min}}^{x^p_{\max}} \mathrm{d}x^p \int_{y^p_{\min}}^{y^p_{\max}} \mathrm{d}y^p \, \mathrm{PSF}\left(x^p, y^p; \mathbf{x}^m(t)\right)$$
$$= \frac{1}{4}\left[\mathbf{erf}\left(\frac{x^p_{\max}-x^m}{\sigma_{z^m}\sqrt{2}}\right) - \mathbf{erf}\left(\frac{x^p_{\min}-x^m}{\sigma_{z^m}\sqrt{2}}\right)\right]\left[\mathbf{erf}\left(\frac{y^p_{\max}-y^m}{\sigma_{z^m}\sqrt{2}}\right) - \mathbf{erf}\left(\frac{y^p_{\min}-y^m}{\sigma_{z^m}\sqrt{2}}\right)\right]. \tag{S.10}$$

Since this area integral has a closed form solution, for simplicity, we denote it as $g^{m,p}\left(\mathbf{x}^m(t)\right)$. The same may not be true of aberrated PSFs [8] which may be treated using the trapezoidal rule as described in the following paragraph and Fig. S.11.

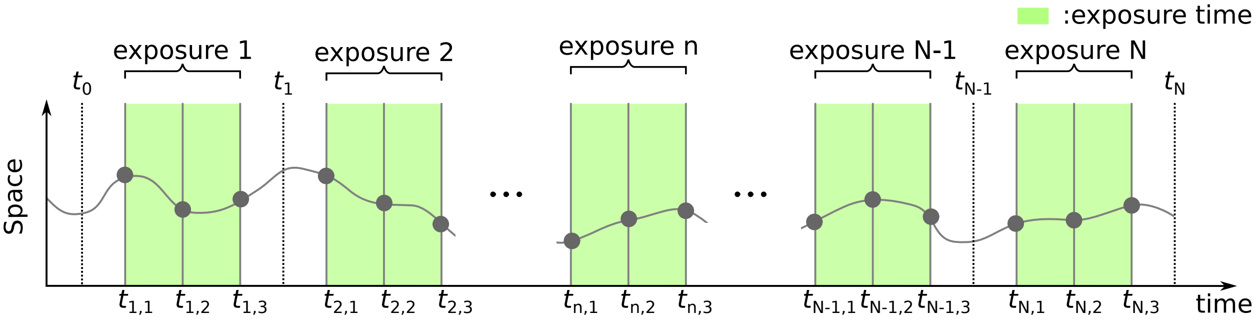

Figure S.11: Discretization schemes for continuous variables $K = 3$.

To clarify, we divide the entire exposure period, denoted as $\tau$, into $K-1$ equal intervals, as depicted in Fig. S.11, with boundaries $t_{n,1}, t_{n,2}, \ldots, t_{n,K}$, such that $t_{n,K} - t_{n,1} = \tau$. We can approximate the integral as follows

$$\int_{t_{n-1}}^{t_n} \mathrm{d}t g^{m,p}\left(\mathbf{x}^m(t)\right) \approx \frac{\delta}{2}\sum_{k=1}^{K-1}\left[g^{m,p}\left(\mathbf{x}^m_{n,k}\right) + g^{m,p}\left(\mathbf{x}^m_{n,k+1}\right)\right] = g^{m,p}_n, \tag{S.11}$$

where $g^{m,p}\left(\mathbf{x}^m_{n,k}\right) \equiv g^{m,p}\left(\mathbf{x}^m\left(t_{n,k}\right)\right)$ and $\delta = \frac{\tau}{K-1}$ is the discretization size. Now Eq. (S.6) becomes

$$u^p_n \approx FA^p\tau + h\sum_{m=1}^{M} b^m g^{m,p}_n. \tag{S.12}$$

From now on, we ignore this last approximation.

#### C.2 Markovian dynamics for spatial trajectories

In Eq. (S.12), the pixel area $A^p$, exposure time $\tau$, background flux $F$, emission rate $h$, and load $b^m$ are input parameters in the data synthesis process. The remaining task is to sample each emitter's spatial trajectory and compute $g^{m,p}_n$. We note that there are two time indices in this model: the frame index $n$ and the within-exposure index $k$, which can make the description cumbersome. To simplify the notation, we define a combined index $l$ as $l = k + (n-1)K$, where $l$ ranges from 1 to $L = NK$. When using this new index, some of the aforementioned variables can be abbreviated as follows

$$t_{(\ell)} \equiv t_{n,k}, \tag{S.13}$$

$$\delta_{(\ell)} \equiv t_{(l+1)} - t_{(\ell)}, \tag{S.14}$$

$$\mathbf{x}^m_{(\ell)} = \left(x^m_{(\ell)}, y^m_{(\ell)}, z^m_{(\ell)}\right) \equiv \left(x^m_{k,n}, y^m_{k,n}, z^m_{k,n}\right). \tag{S.15}$$

It is important to note that we will alternate between the $\ell$ notation and the $n, k$ notation as appropriate in the rest of the text.

In this study, we only consider a single diffusive species. Therefore, the Brownian motion model with a constant diffusion coefficient $D$, can be represented by

$$\mathbf{x}^m_{(\ell)} | \mathbf{x}^m_{(\ell-1)}, D \sim \mathbf{Normal}_3\left(\mathbf{x}^m_{(\ell-1)}, 2D\delta_{(\ell-1)}\mathbf{I}\right). \tag{S.16}$$

Here, the three-dimensional identity matrix $\mathbf{I}$ is included. The initial positions of each emitter are sampled from Normal distributions with fixed means and variances as follows

$$x^m_{(1)} \sim \mathbf{Normal}\left(\mu_{xy}, \sigma^2_{xy}\right), \tag{S.17}$$

$$y^m_{(1)} \sim \mathbf{Normal}\left(\mu_{xy}, \sigma^2_{xy}\right), \tag{S.18}$$

$$z^m_{(1)} \sim \mathbf{Normal}\left(\mu_z, \sigma^2_z\right). \tag{S.19}$$

#### C.3 Model summary

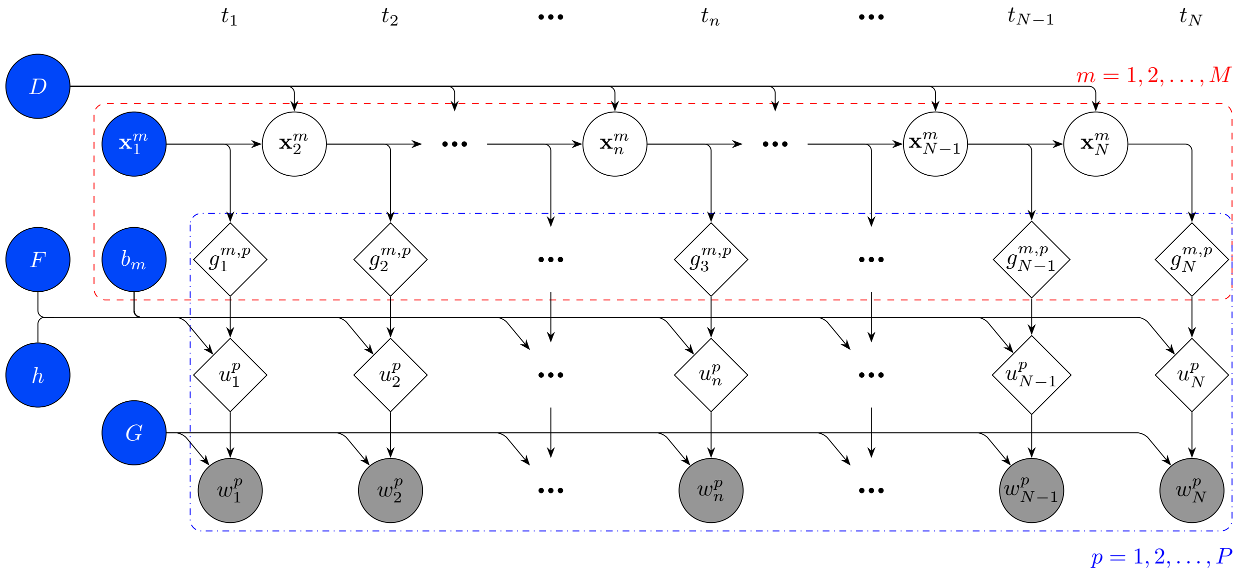

Figure S.12: Graphical representation of our model. Here, circles mark random variables and diamonds mark deterministic variables. Further, image data are shaded.

The entire procedure outlined in Section 3 can be summarized in Fig. S.12. For ease of reference, we have also included a list of all key equations and probability distributions discussed up to this point in the text

$$x^m_{(1)} \sim \textbf{Normal}\left(\mu_{xy}, \sigma^2_{xy}\right), \tag{S.20}$$

$$y^m_{(1)} \sim \textbf{Normal}\left(\mu_{xy}, \sigma^2_{xy}\right), \tag{S.21}$$

$$z^m_{(1)} \sim \textbf{Normal}\left(\mu_{z}, \sigma^2_{z}\right), \tag{S.22}$$

$$\mathbf{x}^m_{(\ell)}|\mathbf{x}^m_{(\ell-1)}, D \sim \textbf{Normal}_3\left(\mathbf{x}^m_{(\ell-1)}, 2D\delta_{(\ell-1)}\mathbf{I}\right) \tag{S.23}$$

$$\begin{aligned} g^{m,p}\left(\mathbf{x}^m_{(\ell)}\right) = \frac{1}{4}&\left[\textbf{erf}\left(\frac{x^p_{\text{max}} - x^m_{(\ell)}}{\sigma_{z^m_{(\ell)}}\sqrt{2}}\right) - \textbf{erf}\left(\frac{x^p_{\text{min}} - x^m_{(\ell)}}{\sigma_{z^m_{(\ell)}}\sqrt{2}}\right)\right] \\ &\times \left[\textbf{erf}\left(\frac{y^p_{\text{max}} - y^m_{(\ell)}}{\sigma_{z^m_{(\ell)}}\sqrt{2}}\right) - \textbf{erf}\left(\frac{y^p_{\text{min}} - y^m_{(\ell)}}{\sigma_{z^m_{(\ell)}}\sqrt{2}}\right)\right], \end{aligned} \tag{S.24}$$

$$g^{m,p}_n = \frac{\delta}{2}\sum_{k=1}^{K-1}\left[g^{m,p}\left(\mathbf{x}^m_{n,k}\right) + g^{m,p}\left(\mathbf{x}^m_{n,k+1}\right)\right], \tag{S.25}$$

$$u^p_n = FA^p\tau + h\sum_{m=1}^{M} b^m g^{m,p}_n, \tag{S.26}$$

$$w^p_n|u^p_n, G \sim \textbf{Gamma}\left(\frac{\beta u^p_n}{f}, fG\right). \tag{S.27}$$

Videos can now be synthesized by following these steps:

1. Specify all the parameters (refer to Tables S.1 and S.3 for details);
2. Sample the initial position of each emitter using Eqs. (S.20) and (S.22);
3. Determine the subsequent spatial trajectory of each emitter using Eq. (S.23);
4. Calculate the photon number $u^p_n$ for each pixel and each frame using Eq. (S.26);
5. Sample from Eq. (S.27) to generate actual measurements.

A note on this data generation is in order. Motion blur artefacts will naturally appear because of step 4 as evaluate Eq. (S.25). Other features such as emitters moving our of focus and their spot becoming broader and blurrier is also incorporated as it follows from Eq. (S.24).

### 4 Inference

The main objective of BNP-Track is to learn spatial trajectories of all emitters, $\mathbf{x}^{1:M}_{(1:L)}$, as well as their loads $b^{1:M}$, diffusion coefficient $D$, background flux $F$, and gain $G$, where $\mathbf{x}^{1:M}_{(1:L)} \equiv \left\{\mathbf{x}^M_{(\ell)}\middle| l = 1, \dots, L, m = 1, \dots, M\right\}$, and $b^{1:M} \equiv \{b^m | m = 1, \dots, M\}$. In the Bayesian paradigm, estimating these quantities is equivalent to sampling from the posterior probability distribution $\mathbb{P}\left(b^{1:M}, \mathbf{x}^{1:M}_{(1:L)}, D, F, G \middle| w^{1:P}_{1:N}\right)$ where $w^{1:P}_{1:N} \equiv \{w^p_n | n = 1, \ldots, N, p = 1, \ldots, P\}$ represents all the measurements. The rest of this section will provide a detailed description of how to sample from this posterior distribution.

#### D.1 Posterior construction

The full posterior probability distribution is written in a specific form and Bayes' theorem is used to derive it

$$\mathbb{P}\left(b^{1:M}, \mathbf{x}^{1:M}_{(1:L)}, D, F, G \middle| w^{1:P}_{1:N}\right) = \frac{\mathbb{P}\left(w^{1:P}_{1:N} \middle| b^{1:M}, \mathbf{x}^{1:M}_{(1:L)}, F, G\right) \mathbb{P}\left(b^{1:M}, \mathbf{x}^{1:M}_{(1:L)}, D, F, G\right)}{\mathbb{P}\left(w^{1:P}_{1:N}\right)}. \quad \text{(S.28)}$$

The probability of $w^{1:P}_{1:N}$ in the denominator, known as the evidence, is a normalization constant and is independent of the variables ($b^{1:M}, \mathbf{x}^{1:M}_{(1:L)}, D, F$, and $G$) that we sample. The probability distribution $\mathbb{P}\left(w^{1:P}_{1:N} \middle| b^{1:M}, \mathbf{x}^{1:M}_{(1:L)}, D, F, G\right)$, called the observation likelihood, can be calculated using Eqs. (S.24) to (S.27). We note that the diffusion coefficient, $D$, does not contribute to the measurements and only the active loads (those with $b^m$ equal to 1) are considered in the calculation of the observation likelihood. Therefore, the observation likelihood can be simplified and denoted as $\mathcal{L}\left(\mathbf{x}^{\mathcal{M}_1}_{(1:L)}, F, G\right)$, which is used frequently throughout the text. The set of all active load positions is denoted as $\mathbf{x}^{\mathcal{M}_1}_{(1:L)}$ with

$$\mathcal{M}_1 = \{m | b^m = 1\}, \; \mathcal{M}_0 = \{m | b^m = 0\}. \quad \text{(S.29)}$$

We also assume that pixels of an EMCCD camera provide independent measurements and measurements from the same pixel are not temporally correlated. This leads to the following equation

$$\mathcal{L}\left(\mathbf{x}^{\mathcal{M}_1}_{(1:L)}, F, G\right) = \prod_{n=1}^{N} \prod_{p=1}^{P} \mathbb{P}\left(w^p_n | b^{1:M}, \mathbf{x}^{1:M}_{(1:L)}, F, G\right). \quad \text{(S.30)}$$

The second term in the numerator of Eq. (S.28), namely the prior, $\mathbb{P}\left(b^{1:M}, \mathbf{x}^{1:M}_{(1:L)}, D, F, G\right)$, can be expanded as follows

$$\mathbb{P}\left(b^{1:M}, \mathbf{x}^{1:M}_{(1:L)}, D, F, G\right) = \mathbb{P}\left(b^{1:M}\right) \mathbb{P}\left(\mathbf{x}^{1:M}_{(1:L)}, D\right) \mathbb{P}\left(F\right) \mathbb{P}\left(G\right). \quad \text{(S.31)}$$

This expansion is valid as the individual probability distributions on the right-hand side are chosen to be independent of each other, as will be detailed in the following sections.

By combining Eqs. (S.28) and (S.31), we obtain the following expression

$$\mathbb{P}\left(b^{1:M}, \mathbf{x}^{1:M}_{(1:L)}, D, F, G \middle| w^{1:P}_{1:N}\right) \propto \mathbb{P}\left(b^{1:M}\right) \mathbb{P}\left(\mathbf{x}^{1:M}_{(1:L)}, D\right) \mathbb{P}\left(F\right) \mathbb{P}\left(G\right) \mathcal{L}\left(\mathbf{x}^{\mathcal{M}_1}_{(1:L)}, F, G\right) \quad \text{(S.32)}$$

which coincides with the posterior probability distribution from which we sample. However, this distribution is too complex to allow for direct sampling. As such, we use a Gibbs sampling scheme to update the parameters iteratively from their own conditional posteriors. Specifically, the sampling procedure follows this sequence

1. Initial values for $\mathbf{x}^{1:M}_{(1:L)}$, $b^{1:M}$, $D$, $F$, and $G$ are set by hand.
2. $G$ is updated by sampling from
$$\mathbb{P}\left(G \middle| w^{1:P}_{1:N}, b^{1:M}, \mathbf{x}^{1:M}_{(1:L)}, F\right) \propto \mathbb{P}\left(G\right) \mathcal{L}\left(\mathbf{x}^{\mathcal{M}_1}_{(1:L)}, F, G\right); \tag{S.33}$$
3. $F$ is updated by sampling from
$$\mathbb{P}\left(F \middle| w^{1:P}_{1:N}, b^{1:M}, \mathbf{x}^{1:M}_{(1:L)}\right) \propto \mathbb{P}\left(F\right) \mathcal{L}\left(\mathbf{x}^{\mathcal{M}_1}_{(1:L)}, F, G\right); \tag{S.34}$$
4. $b^{1:M}$ is updated by sampling from
$$\mathbb{P}\left(b^{1:M} \middle| w^{1:P}_{1:N}, \mathbf{x}^{1:M}_{(1:L)}, F, G\right) \propto \mathbb{P}\left(b^{1:M}\right) \mathcal{L}\left(\mathbf{x}^{\mathcal{M}_1}_{(1:L)}, F, G\right); \tag{S.35}$$
5. $\mathbf{x}^{1:M}_{(1:L)}$ and $D$ are updated by sampling from $\mathbb{P}\left(\mathbf{x}^{1:M}_{(1:L)}, D \middle| w^{1:P}_{1:N}, b^{1:M}, F, G\right)$;
6. $\mathbf{x}^{\mathcal{M}_1}_{(1:L)}$ is updated again by sampling from
$$\mathbb{P}\left(\mathbf{x}^{\mathcal{M}_1}_{(1:L)} \middle| w^{1:P}_{1:N}, D, F, G\right) \propto \mathbb{P}\left(\mathbf{x}^{\mathcal{M}_1}_{(1:L)} \middle| D\right) \mathcal{L}\left(\mathbf{x}^{\mathcal{M}_1}_{(1:L)}, F, G\right). \tag{S.36}$$
7. Repeat step 2 though step 6.

Now, in step 5, we did not expand the conditional posterior as the product of prior and likelihood. This is because it can be decomposed to smaller steps, which are explained in more detail in Section D.5. The other steps are also discussed in Sections D.2 to D.6.

#### D.2 Sampling gain

Our first objective, assuming the camera gain is not pre-calibrated, is to sample camera gain $G$, using Eq. (S.33). To facilitate computation, we select the Inverse Gamma distribution as the prior for $G$, as shown in

$$G \sim \mathbf{InvGamma}\left(\phi_G, \phi_G \chi_G\right). \tag{S.37}$$

By inserting the prior for $G$ from Eq. (S.37) and the camera model from Eq. (S.27) into Eq. (S.33), we obtain

$$\begin{aligned}&\mathbb{P}\left(G \middle| w^{1:P}_{1:N}, b^{1:M}, \mathbf{x}^{1:M}_{(1:L)}, F\right)\\ &= \mathbf{InvGamma}\left(G; \phi_G + \frac{\beta}{f}\sum_{n=1}^{N}\sum_{p=1}^{P} u^p_n, \phi_G\chi_G + \frac{1}{f}\sum_{n=1}^{N}\sum_{p=1}^{P} w^p_n\right)\end{aligned} \tag{S.38}$$

which allows us to directly sample $G$.

#### D.3 Sampling background flux

To sample the background flux $F$, we marginalize over $G$ in Eq. (S.34) by integrating with respect to $G$ as follows

$$\mathbb{P}\left(F \middle| w_{1:N}^{1:P}, b^{1:M}, \mathbf{x}_{(1:L)}^{1:M}\right) = \int dG \mathbb{P}\left(F, G \middle| w_{1:N}^{1:P}, b^{1:M}, \mathbf{x}_{(1:L)}^{1:M}\right). \tag{S.39}$$

By applying Bayes' theorem to the integrand in Eq. (S.39), we obtain

$$\mathbb{P}\left(F \middle| w_{1:N}^{1:P}, b^{1:M}, \mathbf{x}_{(1:L)}^{1:M}\right) \propto \mathbb{P}(F) \int dG \mathbb{P}(G) \mathcal{L}\left(\mathbf{x}_{(1:L)}^{\mathcal{M}_1}, F, G\right). \tag{S.40}$$

The integrand in Eq. (S.40) is proportional to the conditional posterior of $G$ (Eq. (S.33)), which is an Inverse Gamma distribution, as shown in Eq. (S.38). Therefore, this integral can be evaluated as

$$\begin{aligned}&\int dG \mathbb{P}(G) \mathcal{L}\left(\mathbf{x}_{(1:L)}^{\mathcal{M}_1}, F, G\right) \\ &= \left[\prod_{n=1}^{N}\prod_{p=1}^{P} \frac{(w_n^p)^{-1}}{\Gamma\left(\frac{\beta u_n^p}{f}\right)} \left(\frac{w_n^p}{f}\right)^{\frac{\beta u_n^p}{f}}\right] \frac{(\phi_G \chi_G)^{\phi_G}}{\Gamma(\phi_G)} \frac{\Gamma\left(\phi_G + \frac{\beta}{f}\sum_{n=1}^{N}\sum_{p=1}^{P} u_n^p\right)}{\left(\phi_G\chi_G + \frac{1}{f}\sum_{n=1}^{N}\sum_{p=1}^{P} w_n^p\right)^{\phi_G + \frac{\beta}{f}\sum_{n=1}^{N}\sum_{p=1}^{P} u_n^p}}.\end{aligned} \tag{S.41}$$

In our attempt to sample $F$, we can omit all terms in Eq. (S.41) that do not depend on $F$. Therefore, we can express the posterior probability distribution of $F$ as shown in

$$\begin{aligned}&\mathbb{P}\left(F \middle| w_{1:N}^{1:P}, b^{1:M}, \mathbf{x}_{(1:L)}^{1:M}\right) \\ &\propto \mathbb{P}(F) \left[\prod_{n=1}^{N}\prod_{p=1}^{P} \frac{1}{\Gamma\left(\frac{\beta u_n^p}{f}\right)} \left(\frac{w_n^p}{f}\right)^{\frac{\beta u_n^p}{f}}\right] \frac{\Gamma\left(\phi_G + \frac{\beta}{f}\sum_{n=1}^{N}\sum_{p=1}^{P} u_n^p\right)}{\left(\phi_G\chi_G + \frac{1}{f}\sum_{n=1}^{N}\sum_{p=1}^{P} w_n^p\right)^{\phi_G + \frac{\beta}{f}\sum_{n=1}^{N}\sum_{p=1}^{P} u_n^p}},\end{aligned} \tag{S.42}$$

where the gamma function is represented by $\Gamma(\cdot)$. However, no known method exists for directly sampling from Eq. (S.42) regardless of the chosen prior probability of $F$. Therefore, we invoke a Gamma prior on $F$,

$$F \sim \textbf{Gamma}\left(\phi_F, \frac{\psi_F}{\phi_F}\right), \tag{S.43}$$

and apply the Metropolis-Hastings (MH) algorithm to sample from the posterior over $F$. The outline of this process is as follows:

1. Propose a value, $F^{\text{prop}}$, using a proposal distribution based on the current value of $F$ and a proposal distribution, denoted as $F^{\text{old}}$ and $\mathbb{Q}\left(F^{\text{prop}} \middle| F^{\text{old}}\right)$,respectively;

2. Calculate the acceptance ratio $r$ using

$$r = \min\left\{ \frac{\mathbb{P}\left(F^{\text{prop}}\middle|w_{1:N}^{1:P}, b^{1:M}, \mathbf{x}_{(1:L)}^{1:M}\right)}{\mathbb{P}\left(F^{\text{old}}\middle|w_{1:N}^{1:P}, b^{1:M}, \mathbf{x}_{(1:L)}^{1:M}\right)} \frac{\mathbb{Q}\left(F^{\text{old}}\middle|F^{\text{prop}}\right)}{\mathbb{Q}\left(F^{\text{prop}}\middle|F^{\text{old}}\right)}, 1 \right\}; \tag{S.44}$$

3. Generate a uniformly distributed random real number between 0 and 1;

4. If the acceptance ratio is greater than the random number, set $F$ equal to $F^{\text{prop}}$, otherwise set $F$ equal to the old value $F^{\text{old}}$.

To propose a value for $F^{\text{prop}}$, we employ a multiplicative random walker with a 50% probability of setting $F^{\text{prop}}$ to either the product or the quotient of $F^{\text{old}}$ and a value $\epsilon^{\text{prop}}$ drawn from a beta distribution. Part of the acceptance ratio for this process in Eq. (S.44) is given by

$$\frac{\mathbb{Q}\left(F^{\text{old}}\middle|F^{\text{prop}}\right)}{\mathbb{Q}\left(F^{\text{prop}}\middle|F^{\text{old}}\right)} = \frac{F^{\text{prop}}}{F^{\text{old}}} \tag{S.45}$$

which states that the ratio of the probability of the old value given the proposed value to the probability of the proposed value given the old value is equal to the ratio of the proposed value to the old value. In this study, we use the Beta proposal distribution specified in

$$\epsilon^{\text{prop}} \sim \mathbf{Beta}\left(\alpha_\epsilon, 1\right) \tag{S.46}$$

where the parameter $\alpha_\epsilon$ is chosen manually.

Now that all the terms in Eq. (S.44) are specified, we can update $F$ according to the previously outlined four-step procedure. In order to ensure numerical stability in BNP-Track, both Eqs. (S.42) and (S.44) are calculated in logarithmic space. For reference, we provide the complete form of the log acceptance ratio as follows

$$\begin{aligned}
\ln r = &\sum_{n=1}^{N}\sum_{p=1}^{P}\left[\frac{\beta\left(u_n^{p,\text{prop}} - u_n^{p,\text{old}}\right)}{f}\ln\frac{w_n^p}{f} - \ln\Gamma\left(\frac{\beta u_n^{p,\text{prop}}}{f}\right) + \ln\Gamma\left(\frac{\beta u_n^{p,\text{old}}}{f}\right)\right] \\
&+ \ln\Gamma\left(\phi_G + \frac{\beta}{f}\sum_{n=1}^{N}\sum_{p=1}^{P} u_n^{p,\text{prop}}\right) - \ln\Gamma\left(\phi_G + \frac{\beta}{f}\sum_{n=1}^{N}\sum_{p=1}^{P} u_n^{p,\text{old}}\right) \\
&- \left(\frac{\beta}{f}\sum_{n=1}^{N}\sum_{p=1}^{P}\left(u_n^{p,\text{prop}} - u_n^{p,\text{old}}\right)\right)\ln\left(\phi_G\chi_G + \frac{1}{f}\sum_{n=1}^{N}\sum_{p=1}^{P} w_n^p\right) \\
&+ \left(\phi_F - 1\right)\ln\frac{F^{\text{prop}}}{F^{\text{old}}} - \frac{\phi_F}{\psi_F}\left(F^{\text{prop}} - F^{\text{old}}\right).
\end{aligned} \tag{S.47}$$

Here, $u_n^{p,\text{prop}}$ and $u_n^{p,\text{old}}$ represent the values of $u_n^p$ calculated using $F^{\text{prop}}$ and $F^{\text{old}}$, respectively.

#### D.4 Sampling loads

The conditional posterior distribution over the loads, as given in Eq. (S.35), is proportional to the product of the prior over loads and the observation likelihood. As a prior, we assume that the loads over each emitter are independent of each other, which can be expressed as a product of the probability of each load

$$\mathbb{P}\left(b^{1:M}\right) = \prod_{m=1}^{M} \mathbb{P}\left(b^{m}\right). \tag{S.48}$$

Since each load can only be on (1) or off (0), we use a Bernoulli prior for each load,

$$b^{m} \sim \mathbf{Bernoulli}\left(\frac{\gamma}{M}\right) \tag{S.49}$$

with the form chosen such that the expected number of on loads is equal to a specified value $\gamma$. The value of $M$, as it sets the maximum number of emitters to track, should be as large as possible while still being computationally feasible. As a lower limit, $M$ should be at least ten times larger than the value of $\gamma$.

Now we can proceed to update variables $b^1$ through $b^M$ iteratively. The math remains the same for each load, so we will just demonstrate the general case of updating $b^m$. The individual posterior of $b^m$ is given by

$$\mathbb{P}\left(b^{m} \middle| w^{1:P}_{1:N}, \mathbf{x}^{1:M}_{(1:L)}, F, G\right) \propto \mathbb{P}\left(b^{m}\right) \mathcal{L}\left(\mathbf{x}^{\mathcal{M}_1}_{(1:L)}, F, G\right). \tag{S.50}$$

Then the probabilities of both possible values of $b^m$ are

$$\begin{aligned}
\mathbb{P}_{\text{off}} &\equiv \mathbb{P}\left(b^{m} = 0 \middle| b^{1:m-1,m+1:M}, w^{1:P}_{1:N}, \mathbf{x}^{1:M}_{(1:L)}, F, G\right) \\
&= A\frac{M-\gamma}{M}\mathcal{L}\left(b^{1:m-1,m+1:M}, b^{m} = 0, \mathbf{x}^{1:M}_{(1:L)}, F, G\right), && \text{(S.51)} \\
\mathbb{P}_{\text{on}} &\equiv \mathbb{P}\left(b^{m} = 1 \middle| b^{1:m-1,m+1:M}, w^{1:P}_{1:N}, \mathbf{x}^{1:M}_{(1:L)}, F, G\right) \\
&= A\frac{\gamma}{M}\mathcal{L}\left(b^{1:m-1,m+1:M}, b^{m} = 1, \mathbf{x}^{1:M}_{(1:L)}, F, G\right), && \text{(S.52)}
\end{aligned}$$

where $A$ is a normalization constant. Therefore, each $b^m$ is updated according to

$$b^{m} \sim \mathbf{Bernoulli}\left(\mathbb{P}_{\text{on}}\right). \tag{S.53}$$

As in Section D.3, it is important to consider numerical stability when implementing the load sampling scheme. As before, all operations are performed in logarithmic space. Most of the modifications are straightforward algebraic manipulations, but it is worth noting that Eq. (S.53) in logarithmic space can be sampled using the "Gumbel-max" trick [9]. Specifically, let

$$g_{\text{on}} \sim \mathbf{Gumbel}(0,1) \tag{S.54}$$

and

$$g_{\text{off}} \sim \mathbf{Gumbel}(0,1) \tag{S.55}$$

then

$$b^m = \begin{cases} 0, & \text{if } g_{\text{on}} + \ln \mathbb{P}_{\text{on}} < g_{\text{off}} + \ln \mathbb{P}_{\text{off}}; \\ 1, & \text{if } g_{\text{on}} + \ln \mathbb{P}_{\text{on}} \geq g_{\text{off}} + \ln \mathbb{P}_{\text{off}}. \end{cases} \tag{S.56}$$

In this paragraph, we will demonstrate an example of updating two loads, $b^{m_1}$ and $b^{m_2}$ (where $m_2 > m_1$), simultaneously. The majority of the derivation in this section remains valid, and the posterior of $b^{m_1}$ and $b^{m_2}$ becomes a Categorical distribution. The probabilities for each category, $(0,0)$, $(1,0)$, $(0,1)$, and $(1,1)$, are given by

$$\mathbb{P}_{(0,0)} = A\frac{(M-\gamma)^2}{M^2}\mathcal{L}\left(b^{1:m_1-1,m_1+1:m_2-1,m_2+1:M}, b^{m_1}=0, b^{m_2}=0, \mathbf{x}^{1:M}_{(1:L)}, F, G\right), \tag{S.57}$$

$$\mathbb{P}_{(0,1)} = A\frac{\gamma(M-\gamma)}{M^2}\mathcal{L}\left(b^{1:m_1-1,m_1+1:m_2-1,m_2+1:M}, b^{m_1}=0, b^{m_2}=1, \mathbf{x}^{1:M}_{(1:L)}, F, G\right), \tag{S.58}$$

$$\mathbb{P}_{(1,0)} = A\frac{\gamma(M-\gamma)}{M^2}\mathcal{L}\left(b^{1:m_1-1,m_1+1:m_2-1,m_2+1:M}, b^{m_1}=1, b^{m_2}=0, \mathbf{x}^{1:M}_{(1:L)}, F, G\right), \tag{S.59}$$

$$\mathbb{P}_{(1,1)} = A\frac{\gamma^2}{M^2}\mathcal{L}\left(b^{1:m_1-1,m_1+1:m_2-1,m_2+1:M}, b^{m_1}=1, b^{m_2}=1, \mathbf{x}^{1:M}_{(1:L)}, F, G\right). \tag{S.60}$$

The "Gumbel-max" technique should also be modified accordingly in order to accommodate more categories. Specifically, we need to independently sample $g_{(0,0)}$, $g_{(0,1)}$, $g_{(1,0)}$, and $g_{(1,1)}$ and determine the category the maximizes $g + \ln \mathbb{P}$.

#### D.5 Sampling trajectories and diffusion coefficient

As previously mentioned in Section D.1, we know that an emitter only contributes to observations when its load is active ($b^m = 1$). This also implies that the posterior probability distribution of the trajectory of emitters with active loads includes observations, while the distribution of emitters with inactive loads does not. Therefore, to sample from the target distribution $\mathbb{P}\left(\mathbf{x}^{1:M}_{(1:L)}, D \middle| w^{1:P}_{1:N}, b^{1:M}, F, G\right)$, we can update the trajectories of both active ($\mathbf{x}^{\mathcal{M}_1}_{(1:L)}$) and inactive ($\mathbf{x}^{\mathcal{M}_0}_{(1:L)}$) emitters separately. This results in the following posterior form

$$\mathbb{P}\left(\mathbf{x}^{1:M}_{(1:L)}, D \middle| w^{1:P}_{1:N}, b^{1:M}, F, G\right) = \mathbb{P}\left(\mathbf{x}^{\mathcal{M}_0}_{(1:L)} \middle| D\right)\mathbb{P}\left(D \middle| \mathbf{x}^{\mathcal{M}_1}_{(1:L)}\right)\mathbb{P}\left(\mathbf{x}^{\mathcal{M}_1}_{(1:L)} \middle| w^{1:P}_{1:N}, F, G\right). \tag{S.61}$$

Using the logic of ancestral sampling, we can now sample all trajectories and diffusion coefficient in three steps:

1. Sample the trajectories of emitters with active loads from $\mathbb{P}\left(\mathbf{x}^{\mathcal{M}_1}_{(1:L)} \middle| w^{1:P}_{1:N}, F, G\right)$;

2. Sampling the diffusion coefficient from $\mathbb{P}\left(D \middle| \mathbf{x}^{\mathcal{M}_1}_{(1:L)}\right)$;

3. Sample the trajectories of emitters with inactive loads from $\mathbb{P}\left(\mathbf{x}^{\mathcal{M}_0}_{(1:L)} \middle| D\right)$.

##### D.5.1 Active trajectories

The target distribution of active trajectories, denoted by $\mathbb{P}\left(\mathbf{x}_{(1:L)}^{\mathcal{M}_1}\middle|w_{1:N}^{1:P},F,G\right)$, has not yet been derived. Therefore, we must first derive this distribution. We begin by writing

$$\mathbb{P}\left(\mathbf{x}_{(1:L)}^{\mathcal{M}_1}\middle|w_{1:N}^{1:P},F,G\right)=\frac{1}{\mathbb{P}\left(w_{1:N}^{1:P}\right)}\mathbb{P}\left(w_{1:N}^{1:P},\mathbf{x}_{(1:L)}^{\mathcal{M}_1}\middle|w_{1:N}^{1:P},F,G\right). \tag{S.62}$$

After omitting the evidence term (the normalization), this becomes

$$\mathbb{P}\left(\mathbf{x}_{(1:L)}^{\mathcal{M}_1}\middle|w_{1:N}^{1:P},F,G\right)\propto\mathbb{P}\left(w_{1:N}^{1:P},\mathbf{x}_{(1:L)}^{\mathcal{M}_1}\middle|F,G\right). \tag{S.63}$$

Next, similar to the marginalization trick used in Eq. (S.39), we can write

$$\begin{aligned}\mathbb{P}\left(w_{1:N}^{1:P},\mathbf{x}_{(1:L)}^{\mathcal{M}_1}\middle|F,G\right)&\propto\int_0^\infty \mathrm{d}D\int \mathrm{d}\mathbf{x}_{(1:L)}^{\mathcal{M}_0}\mathbb{P}\left(D,w_{1:N}^{1:P},\mathbf{x}_{(1:L)}^{\mathcal{M}_0},\mathbf{x}_{(1:L)}^{\mathcal{M}_1}\middle|F,G\right)\\&\propto\mathcal{L}\left(\mathbf{x}_{(1:L)}^{\mathcal{M}_1},F,G\right)\int_0^\infty \mathrm{d}D\mathbb{P}\left(D\right)\int \mathrm{d}\mathbf{x}_{(1:L)}^{\mathcal{M}_0}\mathbb{P}\left(\mathbf{x}_{(1:L)}^{\mathcal{M}_0},\mathbf{x}_{(1:L)}^{\mathcal{M}_1}\middle|D\right).\end{aligned} \tag{S.64}$$

To evaluate this double integral, we start with writing

$$\mathbb{P}\left(\mathbf{x}_{(1:L)}^{\mathcal{M}_0},\mathbf{x}_{(1:L)}^{\mathcal{M}_1}\middle|D\right)=\mathbb{P}\left(\mathbf{x}_{(1:L)}^{\mathcal{M}_0}\middle|D\right)\mathbb{P}\left(\mathbf{x}_{(1:L)}^{\mathcal{M}_1}\middle|D\right), \tag{S.65}$$

as the trajectories of different emitters are de-correlated. Additionally, $\int \mathrm{d}\mathbf{x}_{(1:L)}^{\mathcal{M}_0}$ represents the integration of every single element of $\mathbf{x}_{(1:L)}^{\mathcal{M}_0}$ over the entire real axis. This leads to

$$\int \mathrm{d}\mathbf{x}_{(1:L)}^{\mathcal{M}_0}\mathbb{P}\left(\mathbf{x}_{(1:L)}^{\mathcal{M}_0},\mathbf{x}_{(1:L)}^{\mathcal{M}_1}\middle|D\right)=\mathbb{P}\left(\mathbf{x}_{(1:L)}^{\mathcal{M}_1}\middle|D\right)\int \mathrm{d}\mathbf{x}_{(1:L)}^{\mathcal{M}_0}\mathbb{P}\left(\mathbf{x}_{(1:L)}^{\mathcal{M}_0}\middle|D\right)=\mathbb{P}\left(\mathbf{x}_{(1:L)}^{\mathcal{M}_1}\middle|D\right), \tag{S.66}$$

which results in

$$\mathbb{P}\left(\mathbf{x}_{(1:L)}^{\mathcal{M}_1}\middle|w_{1:N}^{1:P},F,G\right)\propto\mathcal{L}\left(\mathbf{x}_{(1:L)}^{\mathcal{M}_1},F,G\right)\int_0^\infty \mathrm{d}D\mathbb{P}\left(D\right)\mathbb{P}\left(\mathbf{x}_{(1:L)}^{\mathcal{M}_1}\middle|D\right). \tag{S.67}$$

To continue, we must calculate the probability of active trajectories, $\mathbf{x}_{(1:L)}^{\mathcal{M}_1}$, given the diffusion coefficient, $D$, and specify the prior over $D$, $\mathbb{P}\left(D\right)$. As previously mentioned in Eq. (S.23), we consider the Brownian motion model in this study. Therefore, the probability of $\mathbf{x}(1:L)^{\mathcal{M}1}$ given $D$ can be expressed as

$$\mathbb{P}\left(\mathbf{x}_{(1:L)}^{\mathcal{M}_1}\middle|D\right)=\prod_{m\in\mathcal{M}_1}\left[\prod_{l=1}^{L-1}\mathbb{P}\left(\mathbf{x}_{(l+1)}^{m}\middle|\mathbf{x}_{(\ell)}^{m},D\right)\right]\mathbb{P}\left(\mathbf{x}_{(1)}^{m}\right). \tag{S.68}$$

In this equation, $\mathbb{P}\left(\mathbf{x}_{(1)}^{m}\right)$ is the prior probability distribution for the initial position of the emitter, which is independent of $D$ and therefore constant in the integral of Eq. (S.67). We will address this distribution shortly.

The Brownian motion model also dictates that

$$\mathbb{P}\left(\mathbf{x}^m_{(l+1)}\middle|\mathbf{x}^m_{(\ell)}, D\right) = \mathbf{Normal}_3\left(\mathbf{x}^m_{(l+1)}; \mathbf{x}^m_{(\ell)}, 2D\delta_{(\ell-1)}\mathbf{I}\right)$$
$$\propto D^{-\frac{3}{2}} \exp\left(-\frac{\left|\mathbf{x}^m_{(l+1)} - \mathbf{x}^m_{(\ell)}\right|^2}{4D\delta_{(\ell)}}\right). \tag{S.69}$$

Multiplying the equation above over all emitters with load being 1 and all time points yields

$$\prod_{m\in\mathcal{M}_1}\prod_{l=1}^{L-1}\mathbb{P}\left(\mathbf{x}^m_{(l+1)}\middle|\mathbf{x}^m_{(\ell)}, D\right) \propto D^{-\frac{3}{2}B(L-1)} \exp\left(-\sum_{m\in\mathcal{M}_1}\sum_{l=1}^{L-1}\frac{\left|\mathbf{x}^m_{(l+1)} - \mathbf{x}^m_{(\ell)}\right|^2}{4D\delta_{(\ell)}}\right), \tag{S.70}$$

where $B$ is the cardinality (number of elements) of the active load set $\mathcal{M}_1$. This $D$-dependence can actually be represented by an Inverse Gamma distribution of $D$. Put differently, the Inverse Gamma distribution is the conjugate prior to this likelihood. Therefore, we choose an Inverse Gamma distribution, or conjugate prior, for the diffusion coefficient as

$$D \sim \mathbf{InvGamma}\left(\phi_D, \phi_D\chi_D\right). \tag{S.71}$$

By doing so, we obtain the following equation

$$\mathbb{P}(D)\,\mathbb{P}\left(\mathbf{x}^{\mathcal{M}_1}_{(1:L)}\middle|D\right)$$
$$\propto \left[\prod_{m\in\mathcal{M}_1}\mathbb{P}\left(\mathbf{x}^m_{(1)}\right)\right] D^{-\frac{3}{2}B(L-1)-\phi_D-1} \exp\left(-\frac{\phi_D\chi_D}{D} - \sum_{m\in\mathcal{M}_1}\sum_{l=1}^{L-1}\frac{\left|\mathbf{x}^m_{(l+1)} - \mathbf{x}^m_{(\ell)}\right|^2}{4D\delta_{(\ell)}}\right). \tag{S.72}$$

By substituting Eqs. (S.71) and (S.72) into the integrand of Eq. (S.67) and evaluating the integral, we get

$$\int_0^\infty \mathrm{d}D\mathbb{P}\left(\mathbf{x}^{\mathcal{M}_1}_{(1:L)}\middle|D\right)\mathbb{P}(D)$$
$$\propto \left[\prod_{m\in\mathcal{M}_1}\mathbb{P}\left(\mathbf{x}^m_{(1)}\right)\right]\left(\phi_D\chi_D + \sum_{m\in\mathcal{M}_1}\sum_{l=1}^{L-1}\frac{\left|\mathbf{x}^m_{(l+1)} - \mathbf{x}^m_{(\ell)}\right|^2}{4\delta t_{(\ell)}}\right)^{-\frac{3}{2}B(L-1)-\phi_D}. \tag{S.73}$$

Furthermore, we can see from Eqs. (S.20) to (S.22) that the initial positions of all emitters are not correlated. This can be mathematically expressed as

$$\mathbb{P}\left(\mathbf{x}^{\mathcal{M}_1}_{(1)}\right) = \prod_{m\in\mathcal{M}_1}\mathbf{Normal}\left(x^m_{(1)}; \mu_{xy}, \sigma^2_{xy}\right)\mathbf{Normal}\left(y^m_{(1)}; \mu_{xy}, \sigma^2_{xy}\right)\mathbf{Normal}\left(z^m_{(1)}; \mu_z, \sigma^2_z\right). \tag{S.74}$$

Using this information, we can derive the target distribution from which we can sample trajectories associated with loads 1.

$$\begin{aligned}
&\mathbb{P}\left(\mathbf{x}^{\mathcal{M}_1}_{(1:L)}\middle| w^{1:P}_{1:N}, F, G\right) \\
&\propto \mathbb{P}\left(\mathbf{x}^{\mathcal{M}_1}_{(1)}\right) \mathcal{L}\left(\mathbf{x}^{\mathcal{M}_1}_{(1:L)}, F, G\right) \left(\phi_D\chi_D + \sum_{m\in\mathcal{M}_1}\sum_{l=1}^{L-1} \frac{\left|\mathbf{x}^m_{(l+1)} - \mathbf{x}^m_{(\ell)}\right|^2}{4\delta t_{(\ell)}}\right)^{-\frac{3}{2}B(L-1)-\phi_D} .
\end{aligned} \tag{S.75}$$

##### D.5.2 MH random walk

Since the fully written-out posterior probability distribution in Eq. (S.75) is not suitable for direct sampling, we use the MH algorithm to generate new samples (for what paramaters?). The process is the same as described in Section D.3, with the only changes being the proposal distribution and the acceptance ratio formula.

In the proposal distribution, we add a Normal perturbation to the current trajectory to produce proposed trajectories

$$\mathbb{Q}\left(\mathbf{x}^{\mathcal{M}_1,\text{prop}}_{(1:L)}\middle|\mathbf{x}^{\mathcal{M}_1,\text{old}}_{(1:L)}\right) = \mathbf{Normal}_3\left(\mathbf{x}^{\mathcal{M}_1,\text{prop}}_{(1:L)}; \mathbf{x}^{\mathcal{M}_1,\text{old}}_{(1:L)}, \mathbf{\Sigma_x}\right). \tag{S.76}$$

The perturbation size is controlled by the elements of the diagonal matrix $\mathbf{\Sigma_x}$, which can be manually set or automatically tuned for a target acceptance ratio. This proposal is advantageous because it satisfies the symmetry $\mathbb{Q}\left(\mathbf{x}^{\mathcal{M}_1,\text{prop}}_{(1:L)}\middle|\mathbf{x}^{\mathcal{M}_1,\text{old}}_{(1:L)}\right) = \mathbb{Q}\left(\mathbf{x}^{\mathcal{M}_1,\text{old}}_{(1:L)}\middle|\mathbf{x}^{\mathcal{M}_1,\text{prop}}_{(1:L)}\right)$, which allows us to simplify the acceptance ratio to

$$r = \min\left\{\frac{\mathbb{P}\left(\mathbf{x}^{\mathcal{M}_1,\text{prop}}_{(1:L)}\middle| w^{1:P}_{1:N}, F, G\right)}{\mathbb{P}\left(\mathbf{x}^{\mathcal{M}_1,\text{old}}_{(1:L)}\middle| w^{1:P}_{1:N}, F, G\right)}, 1\right\}, \tag{S.77}$$

Based on Eq. (S.75), the ratio between the posterior probabilities can be expressed as

$$\begin{aligned}
\frac{\mathbb{P}\left(\mathbf{x}^{\mathcal{M}_1,\text{prop}}_{(1:L)}\middle| w^{1:P}_{1:N}, F, G\right)}{\mathbb{P}\left(\mathbf{x}^{\mathcal{M}_1,\text{old}}_{(1:L)}\middle| w^{1:P}_{1:N}, F, G\right)} &= \frac{\mathcal{L}\left(\mathbf{x}^{\mathcal{M}_1,\text{prop}}_{(1:L)}, F, G\right)}{\mathcal{L}\left(\mathbf{x}^{\mathcal{M}_1,\text{old}}_{(1:L)}, F, G\right)} \frac{\mathbb{P}\left(\mathbf{x}^{1:M,\text{prop}}_{(1)}\right)}{\mathbb{P}\left(\mathbf{x}^{1:M,\text{old}}_{(1)}\right)} \\
&\times \left(\frac{\phi_D\chi_D + \sum_{m\in\mathcal{M}_1}\sum_{l=1}^{L-1} \frac{\left|\mathbf{x}^{m,\text{prop}}_{(l+1)} - \mathbf{x}^{m,\text{prop}}_{(\ell)}\right|^2}{4\delta_{(\ell)}}}{\phi_D\chi_D + \sum_{m\in\mathcal{M}_1}\sum_{l=1}^{L-1} \frac{\left|\mathbf{x}^{m,\text{old}}_{(l+1)} - \mathbf{x}^{m,\text{old}}_{(\ell)}\right|^2}{4\delta_{(\ell)}}}\right)^{-\frac{3}{2}B(L-1)-\phi_D} .
\end{aligned} \tag{S.78}$$

As previously mentioned in Section D.3, the acceptance ratio and all probability densities in the implementation of this sampler are calculated in logarithmic space. The terms in Eq. (S.78) have been defined earlier in Eq. (S.30), Eq. (S.77) and Eqs. (S.24) to (S.27), so they will not be further discussed here. Another computational concern when implementing this algorithm is the mixing time, or the time it takes to find the samples with the maximum posterior. In order to achieve optimal performance, it is necessary to find a balance between the acceptance ratio and perturbation size. In particular, this active trajectory sampler allows for partial perturbation, which means that we can perturb segments of one or multiple emitter trajectories in a random order while keeping other parts of the trajectories unchanged. We note that this simply means that elements in the matrix $\mathbf{\Sigma_x}$ from Eq. (S.76) are set to zero, in turn, and this does not affect the mathematical derivations of this section.

##### D.5.3 Sampling diffusion coefficient and inactive trajectories

Similar to the derivations in the previous sections, we begin by defining the target distribution $\mathbb{P}\left(D\middle|\mathbf{x}_{(1:L)}^{\mathcal{M}_1}\right)$, which can be expressed using Bayes' theorem as

$$\mathbb{P}\left(D\middle|\mathbf{x}_{(1:L)}^{\mathcal{M}_1}\right) \propto \mathbb{P}\left(\mathbf{x}_{(1:L)}^{\mathcal{M}_1}\middle|D\right)\mathbb{P}\left(D\right). \tag{S.79}$$

Both terms on the right side of Eq. (S.79) were previously discussed when calculating the integrand in Eq. (S.72). Therefore, for the posterior $\mathbb{P}\left(D\middle|\mathbf{x}_{(1:L)}^{\mathcal{M}_1}\right)$, we write

$$\mathbb{P}\left(D\middle|\mathbf{x}_{(1:L)}^{\mathcal{M}_1}\right) \propto D^{-\frac{3}{2}B(L-1)-\phi_D-1}\exp\left(-\frac{\phi_D\chi_D}{D} - \sum_{m\in\mathcal{M}_1}\sum_{l=1}^{L-1}\frac{\left|\mathbf{x}_{(l+1)}^m - \mathbf{x}_{(\ell)}^m\right|^2}{4D\delta_{(\ell)}}\right), \tag{S.80}$$

or equivalently

$$D \sim \mathbf{InvGamma}\left(\phi_D + \frac{3}{2}B\left(L-1\right), \phi_D\chi_D + \sum_{m\in\mathcal{M}_1}\sum_{l=1}^{L-1}\frac{\left|\mathbf{x}_{(l+1)}^m - \mathbf{x}_{(\ell)}^m\right|^2}{4\delta t_{(\ell)}}\right). \tag{S.81}$$

From this, we can see that the diffusion coefficient can be sampled directly from the resulting posterior probability distribution above.

Finally, we must update the trajectories of emitters with inactive loads. The target distribution in this case is $\mathbb{P}\left(\mathbf{x}_{(1:L)}^{\mathcal{M}_0}\middle|D\right)$. As mentioned at the beginning of Section D.5, these emitters do not contribute to observations and are therefore only sampled based on their prior distribution, defined by Eqs. (S.20) to (S.22) and Eq. (S.23).

#### D.6 Special trajectory proposals

In Section D.5.1, we described how we sample active load trajectories using the MH algorithm with a Normal perturbation proposal distribution. However, because each coordinate of an active load at any time point represents one dimension in the sample space of the target distribution (as shown in Eq. (S.75)), it can be challenging to achieve rapid mixing using only one type of proposal. Therefore, to improve computational efficiency, we have implemented two additional proposal distributions for active load trajectories: the flip proposal and the swap proposal. These "special" proposals take advantage of certain symmetries in our model, while the target distribution remains unchanged.

##### D.6.1 Flip proposal

The flip proposal in our model is derived from the PSF form. As shown in Eqs. (S.7) and (S.8), the considered PSF is symmetric about the focal plane ($xy$-plane). This means that flipping a emitter's spatial position about the focal plane, or mathematically,

$$z_{(\ell)}^{m,\text{prop}} = -z_{(\ell)}^{m},\ z_{(\ell)}^{m,\text{old}} = z_{(\ell)}^{m}, \tag{S.82}$$

does not alter the results of Eqs. (S.24) to (S.27), or the observation likelihood in Eq. (S.30). Similarly, the prior on an emitter's initial $z$-position, as shown in Eq. (S.22), is also symmetric about the focal plane. As a result, the ratio between the target distributions, which is a part of the acceptance ratio, simplifies to

$$\frac{\mathbb{P}\left(\mathbf{x}_{(1:L)}^{\mathcal{M}_1,\text{prop}}\middle|w_{1:N}^{1:P},D,F,G\right)}{\mathbb{P}\left(\mathbf{x}_{(1:L)}^{\mathcal{M}_1,\text{old}}\middle|w_{1:N}^{1:P},D,F,G\right)} = \frac{\mathbb{P}\left(\mathbf{x}_{(1:L)}^{\mathcal{M}_1,\text{prop}}\middle|D\right)}{\mathbb{P}\left(\mathbf{x}_{(1:L)}^{\mathcal{M}_1,\text{old}}\middle|D\right)}. \tag{S.83}$$

Substituting Eq. (S.70) into the previous equation yields

$$\frac{\mathbb{P}\left(\mathbf{x}_{(1:L)}^{\mathcal{M}_1,\text{prop}}\middle|w_{1:N}^{1:P},D,F,G\right)}{\mathbb{P}\left(\mathbf{x}_{(1:L)}^{\mathcal{M}_1,\text{old}}\middle|w_{1:N}^{1:P},D,F,G\right)} = \exp\left(-\sum_{m\in\mathcal{M}_1}\sum_{l=1}^{L-1}\frac{\left|-\mathbf{x}_{(l+1)}^{m,\text{prop}}-\mathbf{x}_{(\ell)}^{m,\text{prop}}\right|^2-\left|\mathbf{x}_{(l+1)}^{m,\text{old}}-\mathbf{x}_{(\ell)}^{m,\text{old}}\right|^2}{4D\delta_{(\ell)}}\right). \tag{S.84}$$

In practice, we randomly select one emitter, denoted as $m'$ and flip all of its positions after a randomly chosen time point $l'$. This allows us to further simplify Eq. (S.84) to

$$\frac{\mathbb{P}\left(\mathbf{x}_{(1:L)}^{\mathcal{M}_1,\text{prop}}\middle|w_{1:N}^{1:P},D,F,G\right)}{\mathbb{P}\left(\mathbf{x}_{(1:L)}^{\mathcal{M}_1,\text{old}}\middle|w_{1:N}^{1:P},D,F,G\right)} = \exp\left(-\frac{\left|-z_{(l'+1)}^{m'}-z_{(l')}^{m'}\right|^2-\left|z_{(l'+1)}^{m'}-z_{(l')}^{m'}\right|^2}{4D\delta_{(\ell)}}\right). \tag{S.85}$$

As flipping an emitter's positions twice should naturally result in the same position, the proposal should therefore be symmetric

$$\frac{\mathbb{Q}\left(\mathbf{x}_{(1:L)}^{\mathcal{M}_1,\text{prop}}\middle|\mathbf{x}_{(1:L)}^{\mathcal{M}_1,\text{old}}\right)}{\mathbb{Q}\left(\mathbf{x}_{(1:L)}^{\mathcal{M}_1,\text{old}}\middle|\mathbf{x}_{(1:L)}^{\mathcal{M}_1,\text{prop}}\right)} = 1. \tag{S.86}$$

The logarithmic acceptance ratio is then given by

$$\ln r = \min\left\{\frac{\left|z_{(l'+1)}^{m'}-z_{(l')}^{m'}\right|^2-\left|-z_{(l'+1)}^{m'}-z_{(l')}^{m'}\right|^2}{4D\delta_{(\ell)}},0\right\}. \tag{S.87}$$

##### D.6.2 Swap proposal

The swapping operation, like the flip proposal, has the advantage of maintaining the invariance of the observation likelihood and being symmetric. To implement this operation, we randomly select $m_1$ and $m_2$, both from the set $\mathcal{M}_1$, and $l = 1, \ldots, L-1$. We then exchange the positions for all subsequent time points according to the following equation

$$\mathbf{x}^{m_1}_{(l:L)} \to \mathbf{x}^{m_2}_{(l:L)},\ \mathbf{x}^{m_2}_{(l:L)} \to \mathbf{x}^{m_1}_{(l:L)}. \tag{S.88}$$

The logarithmic acceptance ratio for this operation is calculated using the following expression

$$\ln r = \min\left\{ \frac{\left|\mathbf{x}^{m_2}_{(l+1)} - \mathbf{x}^{m_2}_{(\ell)}\right|^2 + \left|\mathbf{x}^{m_1}_{(l+1)} - \mathbf{x}^{m_1}_{(\ell)}\right|^2 - \left|\mathbf{x}^{m_2}_{(l+1)} - \mathbf{x}^{m_1}_{(\ell)}\right|^2 - \left|\mathbf{x}^{m_1}_{(l+1)} - \mathbf{x}^{m_2}_{(\ell)}\right|^2}{4D\delta_{(\ell)}}, 0 \right\}. \tag{S.89}$$

#### References

1. Jaqaman, K. *et al.* Robust single-particle tracking in live-cell time-lapse sequences. *Nat. Methods* **5,** 695–702. ISSN: 1548-7105. https://doi.org/10.1038/nmeth.1237 (Aug. 2008).

2. Leary, R. *et al.* Quantitative High-Angle Annular Dark-Field Scanning Transmission Electron Microscope (HAADF-STEM) Tomography and High-Resolution Electron Microscopy of Unsupported Intermetallic GaPd2 Catalysts. *J. Phys. Chem. C* **116,** 13343–13352. eprint: https://doi.org/10.1021/jp212456z. https://doi.org/10.1021/jp212456z (2012).

3. Siepmann, M., Schmitz, G., Bzyl, J., Palmowski, M. & Kiessling, F. *Imaging tumor vascularity by tracing single microbubbles* in *2011 IEEE International Ultrasonics Symposium* (2011), 1906–1909.

4. Hajj, B., El Beheiry, M., Izeddin, I., Darzacq, X. & Dahan, M. Accessing the third dimension in localization-based super-resolution microscopy. *Phys. Chem. Chem. Phys.* **16,** 16340–16348 (2014).

5. He, H. *et al.* Super-Resolution Monitoring of Mitochondrial Dynamics upon Time-Gated Photo-Triggered Release of Nitric Oxide. *Anal. Chem.* **90.** PMID: 29316789, 2164–2169. eprint: https://doi.org/10.1021/acs.analchem.7b04510. https://doi.org/10.1021/acs.analchem.7b04510 (2018).

6. Zhang, B., Zerubia, J. & Olivo-Marin, J.-C. Gaussian approximations of fluorescence microscope point-spread function models. *Appl. Opt.* **46,** 1819–1829 (2007).

7. Sheppard, C. & Matthews, H. Imaging in high-aperture optical systems. *Josa a* **4,** 1354–1360 (1987).

8. Goodman, J. W. *Introduction to Fourier optics* (Roberts and Company publishers, 2005).

9. Jang, E., Gu, S. & Poole, B. *Categorical reparametrization with gumble-softmax* in *International Conference on Learning Representations (ICLR 2017)* (2017).
